# Low protein intake compromises the recovery of lactation-induced bone loss in female mouse dams without affecting skeletal muscles

**DOI:** 10.1101/2020.05.02.073759

**Authors:** Ioannis Kanakis, Moussira Alameddine, Mattia Scalabrin, Rob J. van ‘t Hof, Triantafillos Liloglou, Susan E. Ozanne, Katarzyna Goljanek-Whysall, Aphrodite Vasilaki

## Abstract

Lactation-induced bone loss occurs due to high calcium requirements for fetal growth but skeletal recovery is normally achieved promptly post-weaning. Dietary protein is vital for fetus and mother but the effects of protein undernutrition on the maternal skeleton and skeletal muscles is largely unknown. We used mouse dams fed with normal (N, 20%) or low (L, 8%) protein diet during gestation and lactation and maintained on the same diets (NN, LL) or switched from low to normal (LN) during a 28d skeletal restoration period post lactation. Skeletal muscle morphology and neuromuscular junction integrity was not different between any of the groups. However, dams fed the low protein diet showed extensive bone loss by the end of lactation, followed by full skeletal recovery in NN dams, partial recovery in LN and poor bone recovery in LL dams. Primary osteoblasts from low protein diet fed mice showed decreased *in vitro* bone formation and decreased osteogenic marker gene expression; promoter methylation analysis by pyrosequencing showed no differences in *Bmpr1a, Ptch1, Sirt1, Osx* and *Igf1r* osteoregulators, while miR-26a, -34a and -125b expression was found altered in low protein fed mice. Therefore, normal protein diet is indispensable for maternal musculoskeletal health during the reproductive period.

## INTRODUCTION

Gestation and lactation are two of the most metabolically challenging life phases in females, affecting almost all the physiological systems in the human body. Bone loss arises very rapidly as a result of substantial changes in the maternal skeleton (1). During pregnancy, fetal requirements for calcium are met by a 2-fold up-regulation of intestinal absorption and is thought to be the major mechanism of maternal adaptation, partly mediated by the action of 1,25-vitamin D on intestinal cells (2). This environment allows the maternal skeleton to accumulate sufficient calcium to support bone mineralisation in the fetus during the third trimester (3). Maternal bone mass declines during lactation, when skeletal calcium is released to the breast milk. Although renal calcium excretion is reduced with increased tubular reabsorption (1), this is not sufficient to prevent bone loss. Increased circulating parathyroid hormone related peptide (PTHrP), produced by the lactating breast, plays a key role in calcium release and in combination with low estradiol levels leads to high bone resorption rates (4).

Breast-feeding milk from lactating dams is the main source of food for the neonates and the maternal skeleton provides a large proportion of calcium that is vital for the growing newborn skeletal system (5, 6). The overall bone turnover is increased during lactation, but bone resorption exceeds formation and this leads to substantial bone mass reduction (7, 8). Studies in humans have shown that women’s bone loss can reach around 7%, while this percentage in female mice is greater (up to 30%) due to higher numbers of offspring being suckled during the 3-week lactation period (9, 10). However, weaning triggers skeletal recovery that occurs very rapidly after the end of lactation (10, 11). Bone mass regain is mediated by increased apoptosis of resorptive osteoclasts and increased rates of bone formation (10, 11) with partial or full restoration of mechanical properties (12). Although epidemiological studies in humans suggest that the number of pregnancies has no long-term effects on fracture risk (13, 14), data for the duration of lactation are controversial and some studies report that there is a possible correlation with lower bone mineral density (BMD) in later life (15, 16). It has also been suggested that BMD is fully recovered at the lumbar spine but only partially at the hip in humans (17, 18) and in the tibia in rodents (19). However, the detailed mechanisms of post-weaning maternal bone recovery are largely unknown.

Protein metabolism is also challenging especially during lactation, when nutrient enriched milk production has to be sufficient for the newborn. Protein catabolic rates are increased dramatically and maternal body protein reservoirs, such as the skeletal muscles, are recruited to meet these requirements (20). Protein mobilisation from skeletal muscle is triggered as an adaptive response to enhance protein content in milk through proteolysis and this process is in fine balance with maternal dietary protein intake (21, 22). Muscle metabolic events, such as fatty acid oxidation, are also linked to food intake via nervous system mediated regulation (22). To our knowledge, while muscle proteolysis has been largely studied during lactation, there is no evidence for maternal skeletal muscle fibre morphological changes or neuromuscular junctions (NMJs), the synapse between a motor neuron and a muscle fibre, in conjunction with protein under-nutrition.

Dietary calcium (Ca) and vitamin D supplements during gestation and lactation have been extensively studied, suggesting beneficial effects on both maternal and fetal/offspring skeletal homeostasis (23). Ca supplementation in the maternal diet has been proved beneficial for both maternal and offspring bone health (24); the National Academy of Sciences recommends 1000mg of daily Ca consumption for pregnant and breastfeeding women. Although some studies have explored the consequences of other nutrients, such as soy isoflavone (25) and prebiotics (26) on the maternal skeleton and other organs during lactation and recovery, very little is known about the effects of the maternal protein intake.

In recent years, there has been increasing evidence of the epigenetic regulation of bone health. Epigenetic mechanisms are potential therapeutic targets due to their reversible nature. Several studies have suggested that methylome changes play an important role in osteoblast differentiation and activity (27–30). Hypomethylation of the promoters of the *runt-related transcription factor-2* (*Runx-2*), *bone gamma-carboxyglutamate protein* (*Bglap,* the coding gene for osteocalcin) and *osterix* (*Osx*) genes is involved in osteogenic differentiation of adipose-derived mesenchymal stromal cells (MSCs) (31). Other regulatory mechanisms involve microRNAs (miRs) that can regulate post-transcriptional gene expression. MiRs have been shown to control bone-related genes (32–34). For example, miR-204/211 levels are increased, suppressing *Runx-2* gene expression and promoting adipogenic over osteogenic differentiation of mesenchymal stromal cells (MSCs) (34), while miR-15b induces osteoblast differentiation by inhibiting Runx-2 degradation (35). Moreover, overexpression of miR-2861 and miR-3960 promotes BMP2-induced osteoblastogenesis, and their suppression inhibits osteoblast differentiation (36). It is also known that miRs are essential for endochondral ossification since osteoblast-specific Dicer knockout mice have deficient cortical bone formation and bone integrity (37, 38).

The aim of this study was to investigate the effects of protein restriction on the maternal musculoskeletal system in mouse dams during gestation and lactation as well as the post-weaning recovery period. Histological structural measurements in skeletal muscle and NMJ morphology evaluations were performed. Bone microarchitecture and turnover were assessed using micro computed tomography (microCT) and bone histomorphometry. To delineate the regulatory mechanisms, we also performed *in vitro* experiments to determine potential epigenetic effects on primary bone cells and expression patterns of selected miRs.

## MATERIALS AND METHODS

### Animals

This study included 39 female mice. We used B6.Cg-Tg(Thy1-YFP)16Jrs/J mice, which express yellow fluorescent protein (YFP) only in neuronal cells (Jackson Laboratory; stock number 003709). Mice were housed in individually vented cages maintained at 21 ± 2°C on a 12-hour light/dark cycle. All experimental protocols were performed in compliance with the UK Animals (Scientific Procedures) Act 1986 regulations for the handling and use of laboratory animals and received ethical approval from the University of Liverpool Animal Welfare Ethical Review Committee (AWERB). Mice were monitored daily for any health and welfare issues.

### Experimental groups

All mice were fed *ad libitum* food and water. Solid food pellets comprised of normal protein diet (N, 20% crude protein; Special Diet Services, UK; code 824226) or low protein diet (L, 8% crude protein; Special Diet Services, UK; code 824248) with isocaloric value. Groups of 8 weeks old nulliparous female mice were fed on either L (low) or N (control) protein diet for 2 weeks prior to mating. Once adapted to the diets, the mice were mated with age-matched males on N diet and the pregnant mice were kept on the same diet throughout gestation (19-21 days) and lactation (21 days). Suckling pup number was kept the same for all animals during lactation (n=5-6 pups) to prevent confounding effects of differences in littersize. Following lactation, mice in the normal (group Lac-N, n=8) and low (group Lac-L, n=8) groups were culled (16 weeks old) and tissues were harvested (skeletal muscles and bones). The recovery (Rec) groups were comprised of dams (20 weeks old) fed on N until the end of lactation and remained on the same diet for the recovery (28 days) period (group Rec-NN, n=5), and mice on L diet until weaning which remained on the same diet for recovery (group Rec-LL, n=5) or switched post-weaning to N protein diet (group Rec-LN, n=5) for the recovery period (Fig. 1A). At the end of lactation or recovery mice were euthanised by a rising concentration of CO_2_. Age-matched virgin mice (18 weeks old) were used as a control (group Virgins, n=8). To reduce the number of animals according to 3Rs recommendations, the end-point age of Virgins was kept at the middle of the recovery period (2 weeks after lactation and 2 weeks before recovery end) to serve as controls for both experimental periods (Fig. 1A). At least for the skeletal system, no significant change has been observed in bone mass and structure of female mice between months 3-4 of age (39).

**Figure 1.**
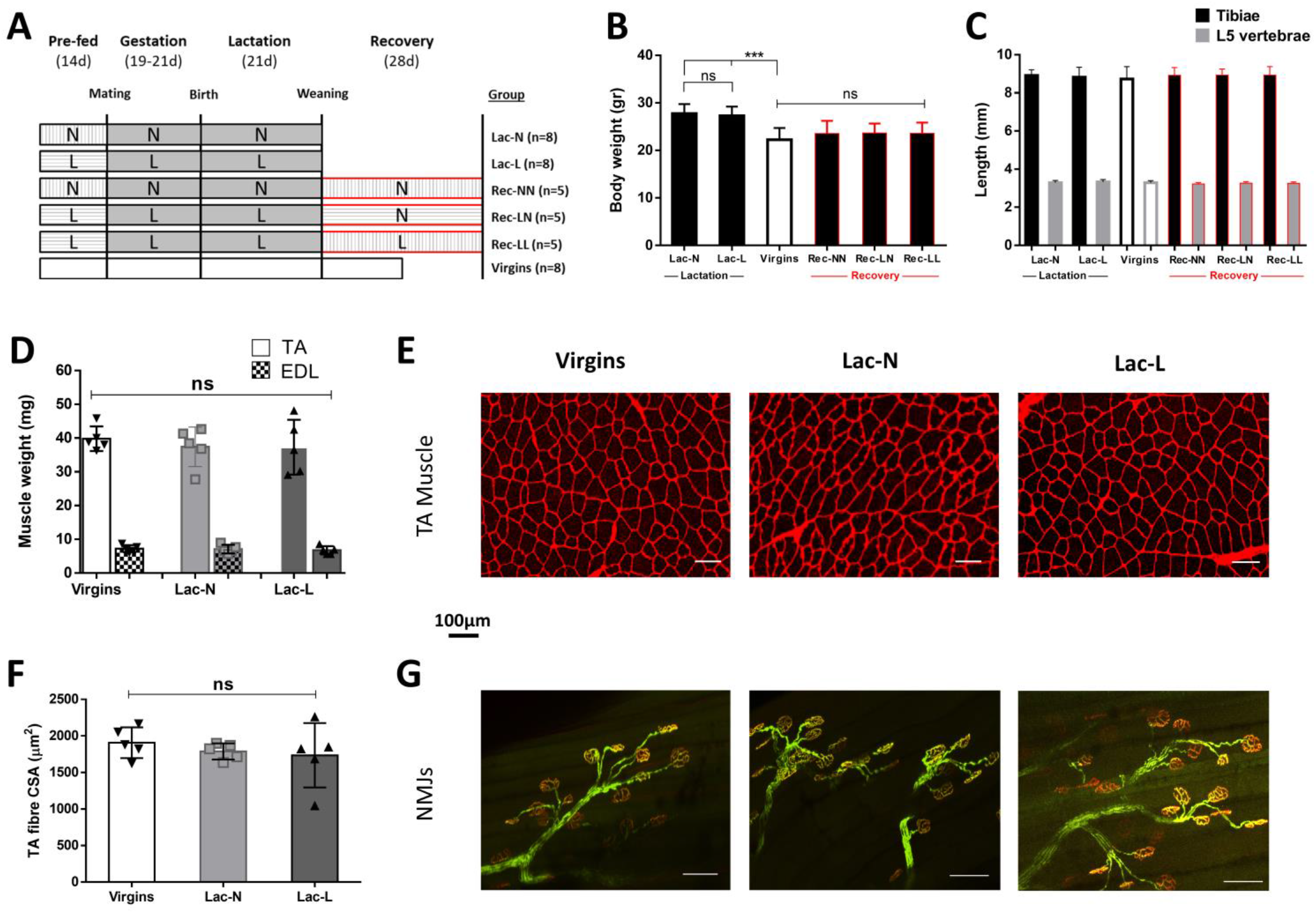
Experimental design of the study (A). Total body weights of the mouse dams used in all experimental groups (B) and the corresponding tibial and L5 vertebral length as measured by microCT showing no difference between the groups (C). The net weights of TA and EDL skeletal muscles were similar in Lac-N (n=8) and Lac-L (n=8) mice when compared to Virgins control (D) and image analysis of transverse cryosections (E) showed no changes in the fibre morphology and fibre CSA in the TA muscles (F). Similarly, NMJs appeared unaffected with perfect overlapping of the pre- (green) and post-synaptic (red) terminals (G). All data are presented as mean±SD. ns: not significant; \*\*\**p*<0.001; scale bar: 100μm.

### Muscle histology and NMJ imaging

#### Skeletal Muscles

Immediately after culling, the extensor digitorum longus (EDL) and tibialis anterior (TA) skeletal muscles were carefully dissected and weighed (n=5). The TA muscles were embedded in Cryomatrix (Thermo Fischer, UK), immediately immersed in liquid nitrogen-frozen isopentane, and stored at −80°C for cryosectioning. TA muscles were placed at −20°C for at least 30 min prior to cryosectioning. Transverse sections (10 μm) were cut using a Leica cryotome and collected on Superfrost glass slides (ThermoScientific, UK). Sections were washed with PBS for 10 min before staining with 1:1000 dilution of rhodamine wheat germ agglutinin (WGA; 5 μg/mL;Vector Laboratories,UK) for 10 min, mounted in antifade medium with DAPI (Vector Laboratories, CA, USA) and were visualised with an Axio Scan.Z1 slide scanner (Zeiss, UK).

The EDL muscles were fixed in 10% neutral buffered formalin (NBF) and stored in PBS with sodium azide at 4°C for NMJs morphological image analysis. Before staining, samples were washed in PBS for 5 minutes and then stained with α-Bungarotoxin-Alexa Fluor™ 647 conjugate (Invitrogen, UK) diluted 1:500 in PBS for at least 30 minutes in a dark environment at room temperature. Muscles were kept in a dark environment until analysis on a Nikon C1 Eclipse Ti confocal laser scanning microscope (Nikon, UK).

#### Bones

The spines and left or right hindlimbs were dissected, cleaned from muscles, fixed in 10% NBF solution for 24 h, extensively washed with PBS and scanned with microCT, followed by decalcification in 10% EDTA for 3 weeks and subsequent storage in 70% EtOH until processing (n=5-8). The other hindlimb was used for primary osteoblasts isolation and cell cultures. Decalcified spines were embedded in paraffin or cryopreserved and embedded in Cryomatrix and sectioned coronally (5-8μm thickness). H&E staining was performed using standard protocol and tartrate-resistant acid phosphatase (TRAP) as previously described (40). Osteoblasts were identified by their cuboidal shape and proximity to endosteal surfaces and osteoclasts by their red colour after TRAP staining. For IHC experiments, cryosections were washed with PBS, blocked with 10% normal goat serum (NGS) and non-immune rabbit serum served as a negative control. Rabbit anti-mouse osteocalcin (Ocn) polyclonal antibody (Abcam, UK, ab39876) was used in 1:500 dilution in 5% NGS to identify osteoblasts. Primary antibodies were detected using a Vectastain ABC kit with a secondary goat anti-rabbit biotinylated antibody and visualized with HRP-conjugated streptavidin using 3,3′-diaminobenzidine (DAB; Vectorlabs, UK). Histomorphometric analyses were performed according to ASBMR standards using open-source software (40, 41).

### Micro-computed tomography (microCT)

Hindlimbs and L5 lumbar vertebra were scanned using a Skyscan 1272 scanner (Bruker, Belgium; 0.5 aluminium filter, 50 kV, 200 mA, voxel size 4.60 μm, 0.3° rotation angle step). Datasets were reconstructed using NRecon and 3D volumes of interest (VOI) were selected using Dataviewer and CTan software (Bruker, Belgium). Trabecular bone parameters were analysed using CTAn in the proximal tibial metaphysis and the vertebral body of L5 vertebra. Cortical bone was analysed at the tibial midshaft. For trabecular bone analysis, VOI was selected using mineralised cartilage as a reference point. The tibial VOI analysed was 400 slices starting 20 levels distal to the reference point, while for cortical bone measurements, a VOI (100 slices) was selected 600 slices below the reference point, as previously described (42, 43). Trabecular bone was automatically separated from cortical bone using a macro in CTAn.

### *In vitro* bone cell culture and mineralisation assay

For the primary bone cell cultures, immediately after microCT scans (<2h from sacrifice), midshafts of long bones were isolated (n=3/group), surrounding muscles removed, and the bones centrifuged (3min at 800×g) to remove the bone marrow. The bone shafts were cut into small pieces using a scalpel. and adhering cells were removed by digestion with collagenase type I (Sigma, 1mg/ml in Hank’s balanced salt solution, HBSS) for 45min in a shaking water bath at 37°C, washed in PBS and cultured in alpha-MEM with Glutamax™ (Gibco, UK) and nucleosides, containing 10% heat-inactivated FBS and penicillin (100 IU/ml)/streptomycin (100 μg/ml) (Invitrogen) in a humidified 5% CO_2_ incubator at 37°C, as previously described (42). Upon reaching semi-confluence, primary osteoblasts (Obs), grown out of the cleaned bone chips, were harvested using trypsin/EDTA (Gibco, UK) and seeded onto 6-well plates (10^5^ cells/well) in osteogenic medium (50μg/mL L-ascorbic acid-2-phosphate and 5mM β-glycerophosphate) (Sigma, UK) for 24 days. Mineralisation capacity was assessed by Alizarin Red S (ARS) (Sigma, UK) staining. Bone nodule surface area was calculated using ImageJ (NIH), as previously described (42, 44).

### Quantitative polymerase chain reaction (qPCR)

To assess gene expression levels, total RNA was isolated from primary osteoblasts using TRiZOL reagent (Invitrogen, CA, USA), cleaned-up using the RNeasy kit (Qiagen, UK) and cDNA was synthesized with the High Capacity cDNA Trancription kit (Applied Biosystems, UK). Expression levels of alkaline phosphatase (*Alp*), collagen type 1 (*Col1a1*) and runt-related transcription factor-2 (*Runx-2*) mRNA were used as osteogenic markers (42) (Supplementary Table 1). qPCR was performed on a RotorGene*™* 6000 (Corbett Research) instrument with SYGR (Bioline, UK) and results were analysed using beta-actin (*Actb*) as a stable reference gene for osteoblasts (45). For miR expression analysis, total RNA (n=3/group) was isolated and purified using the mirVana™ kit (Thermo Fischer, UK), reverse transcription of total RNA containing miRs was performed with miScript II RT kit (Qiagen, UK) and specific primers for miR-26a, 34a and −125b were used for the qPCR utilizing RNU6 as the reference gene. Results were analysed using the modified delta CT method (46) or presented as fold difference as compared to the reference gene levels.

### miR:target prediction and bioinformatics

To predict the targets of the differentially expressed miRs, we used the miRWalk on-line tool (47) by applying simultaneous search from four different databases, including miRWalk, TargetScan (48), miRDB (49) and MiRTarBase (50) using the default parameters of 7 as the minimum seed length at the 3’-UTR site and showing only the statistically significant mRNAs. A total of 174 target genes were obtained. Cytoscape v3.7.2 (51) software was used to build the interaction networks between predicted targets and miRs as well as to determine the biological roles of the target mRNAs utilising Gene Ontology (GO) terms of biological process and molecular functions. The enriched GO terms were presented as enrichment scores. KEGG pathway analysis (52) was performed to determine the involvement of the predicted mRNAs targets in different biological pathways. P<0.05 was considered to indicate a statistically significant result.

#### Promoter methylation analysis

Pyrosequencing methylation assays for bone morphogenetic protein receptor 1a (*Bmpr1*a), patched 1 (*Ptch1*), osterix (*Osx*), insulin-like growth factor 1 receptor (*Igf1r*) and sirtuin 1 (*Sirt1*) gene promoters were designed using the Pyromark Assay Deisgn 2.0 software (Qiagen). PCR and sequencing primers are provided in Supplementary Table 1. Primary osteoblasts-derived genomic DNA (1 μg each) from all the experimental groups (n=3/group) was subjected to bisulphite treatment using the EZ DNA methylation™ kit (Zymo Research, USA) according to the manufacturer’s protocol. Pyrosequencing templates were prepared by PCR amplification (45 cycles) of approximately 30 ng bisulphite-treated DNA using HotStarTaq Master Mix (Qiagen), 150 nM biotinylated primer and 300 nM non-biotinylated primer (Supplementary Table 1). Optimized annealing temperatures were 52°C for *Bmpr1a*, 51°C for *Ptch1*, 48°C for *Sirt1*, 50°C for *Osx* and 55°C for *Igf1r*. PCR efficiency and specificity were verified by agarose gel electrophoresis. PCR products were immobilised on streptavidin coated sepharose beads, and pyrosequenced on PyroMark™Q96 MD instrument (Qiagen) according to the manufacturer’s instructions. The sequence runs were analysed using the Pyromark Q962.5.8 software Q-CpG software.

### Statistical analysis

All data were analysed with GraphPad Prism 6 software and expressed as the mean ± SD. Data sets were tested for normal distribution with the D’Agostino-Pearson normality test. Comparisons between four groups were performed by one-way analysis of variance (ANOVA) followed by Tukey’s multiple comparisons post hoc test where appropriate. For comparisons between two groups, unpaired Student’s t-test or Mann-Whitney U test was applied. In all cases, *P* values less than 0.05 were considered statistically significant.

## RESULTS

### Body weights and bone length

Total body weights were recorded after euthanasia. There was no difference between groups Lac-N and Lac-L (16 weeks old) while weight was increased in both these groups compared to the Virgins group (18 weeks old), as expected (Fig. 1B). For the recovery period, similar weights were observed for all groups of animals (Rec-NN, Rec-LN and Rec-LL) (Fig. 1B). Additionally, there was no difference in tibial and L5 vertebral lengths, as measured by microCT (Fig. 1C).

### Low protein diet has no effect on skeletal muscle and NMJ morphology in mouse dams

We first examined the effects of low protein diet during pregnancy and lactation on skeletal muscle and NMJ morphology. There was no difference in TA and EDL weights between the Virgins, Lan-N and Lac-L groups (Fig. 1D). Furthermore, cross-sectional area (CSA) of the TA skeletal muscle fibres, measured using transverse cryosections, was also similar between groups (Fig. E and F). Detailed confocal image analysis of the EDL muscle also indicated that NMJ morphology was not affected by the reproduction process or the protein content of the diet, as the pre-synaptic YFP-labelled motor axons and the post-synaptic bungorotoxin-stained AchRs perfectly overlapped without any evidence of denervation or clustering (Fig. 1G). In light of these findings, we decided not to proceed with muscle and NMJ analyses for the recovery groups.

### Protein under-nutrition enhances lactation-induced bone loss and delays recovery

We evaluated the effect of low protein intake on tibial and L5 vertebral bone mass by measuring the trabecular and cortical parameters obtained from microCT scans. Our aim was to first examine if protein under-nutrition during gestation and lactation had an impact on skeletal mass and structure and if so, how this diet would affect bone recovery. Therefore, we compared the trabecular %BV/TV in tibiae between Lac-N and Lac-L dams, without using controls as the lactation-induced bone loss has been extensively been reported in rodents (10, 11). Lac-L dams had a significant reduced bone volume in comparison with lactating Lac-N mice on a normal diet, due to decreased trabecular thickness, and decreased trabecular number as well as decreased connectivity and a more rod-like appearance of the trabecular bone (Fig. 2A and G, Supplementary Table 2), reflecting a compromised bone microarchitecture. Similar outcomes were obtained for the trabecular network in L5 vertebral body, showing a further bone mass reduction of 22.75% in Lac-L as compared to Lac-N mice (Fig. 2E and K, Supplementary Table 2). Cortical bone analyses revealed a significant decrease in the cortical thickness in the midshaft of long-bones in lactating mice with greater thickness loss in the Lac-L group, in comparison with Lac-N (Fig. 2C and H).

**Figure 2.**
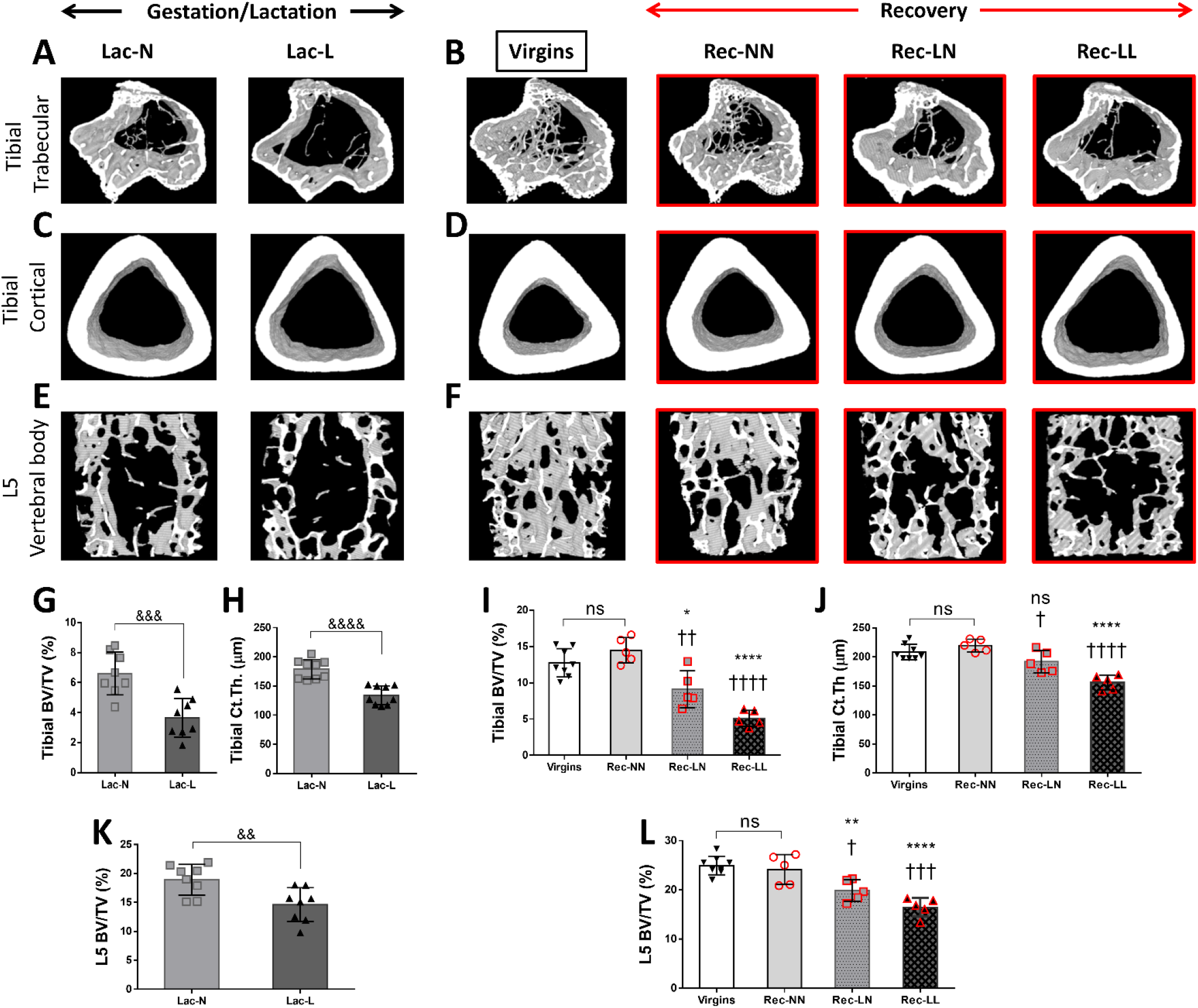
Representative images and quantitative comparisons in Lac-N (n=8) and Lac-L (n=8) mouse dams during gestation/lactation for tibial trabeculae %BVTV (A and G), cortical thickness (C and H) and the L5 trabecular %BVTV (E and K). Tibiae and spines of Rec-NN, Rec-LN and Rec-LL mice (n=5/group) were collected after the 28d recovery and compared to the control Virgins (n=8) group (B and I for tibial trabecular %BVTV; D and J for cortical thickness; F and L for L5 trabecular %BVTV). All data are presented as mean±SD. ns: not significant; ^&&^*p*<0.01, ^&&&^*p*<0.001, ^&&&&^*p*<0.0001 in Lac-N versus Lac-L comparisons. Asterisks indicate comparisons versus Virgins and crosses versus Rec-NN.

After weaning, female mice in the Rec-NN group, which were fed a normal protein diet during gestation and lactation as well as for the recovery period of 28d, showed full recovery of trabecular bone volume and architecture in tibiae (Fig. 2B and I, Supplementary Table 2) and in L5 vertebrae (Fig. 2F and L, Supplementary Table 2), when compared to the Virgins group. Rec-NN mice had statistically significant higher trabecular bone volume than Rec-LN and Rec-LL mice, at both skeletal sites. Cortical bone thickness showed full recovery in the Rec-NN group, partial recovery in the Rec-LN and poor recovery in the Rec-LL (Fig. 2D and J).

### Low protein intake has detrimental effects on bone remodelling

In order to evaluate bone turnover in the spines of mouse dams, we performed bone histomorphometric analyses, as previously described (40). Routine H&E staining (Supplementary Figure 1), confirmed by Ocn IHC (Fig. 3A), showed that Lac-L lactating dams had lower numbers of osteoblasts per bone perimeter (NOb/Bm) and reduced osteoblastic surface per bone surface (ObS/BS) as compared to the Lac-N group (Fig. 3B and C). In the recovery period, Rec-NN mice showed increased osteoblast numbers, in comparison with the Virgin control mice. In contrast, osteoblastic number and surface was lower in the Rec-LN mice while the Rec-LL mice showed no signs of recovery with significantly reduced osteoblast number and surface (Fig. 3D, E and F).

**Figure 3.**
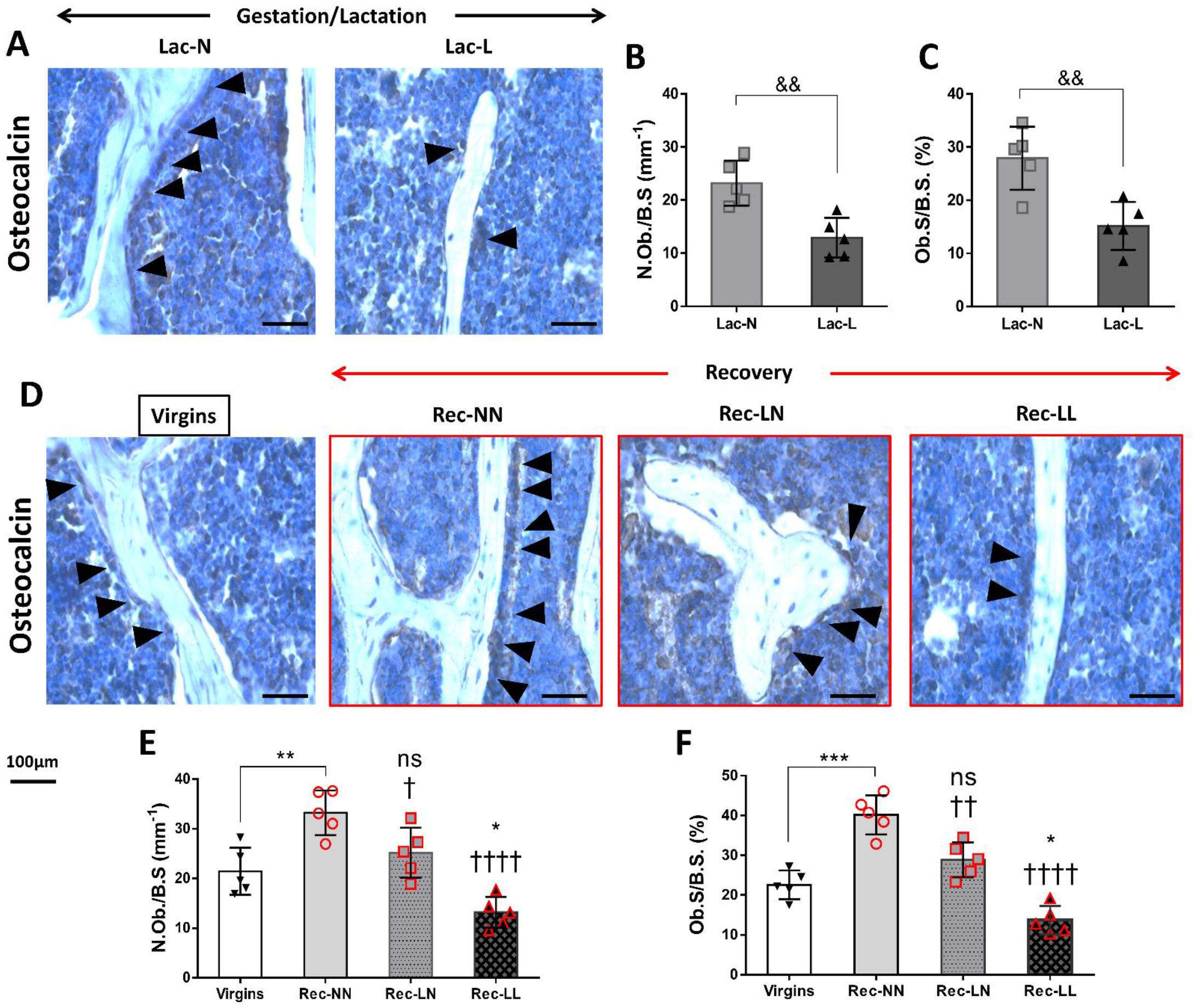
Representative Ocn (A) immunostained L5 vertebral body sections and quantitative comparisons in Lac-N (n=5) and Lac-L (n=5) mouse dams during gestation/lactation for number of osteoblasts per bone surface (N.Ob./B.S.) (B) and the percentage of osteoblast surface per bone surface (Ob.S./B.S.) (C). The corresponding histological images and parameters for the recovery period in Rec-NN (n=5), Rec-LN (n=5) and Rec-LL (n=5) groups are shown in D, E and F, respectively, and were compared to the Virgins control. Arrows indicate Ocn-positive osteoblasts attached to the bone surface. All data are presented as mean±SD. ns: not significant; ^&^*p*<0.05, ^&&^*p*<0.01 in Lac-N versus Lac-L comparisons. Asterisks indicate comparisons versus Virgins and crosses versus Rec-NN. \**p*<0.05, \*\**p*<0.01, \*\*\**p*<0.001, \*\*\*\**p*<0.0001.

TRAcP staining revealed that lactating mice on the low protein diet had increased osteoclast numbers and size compared to the Lac-N group (Fig. 4A). Osteoclast number and size were also increased in the Rec-NN dams compared to the Virgin controls (Fig. 4B and C), indicating a high rate of bone remodelling, most likely coupled with increased bone formation. Interestingly, Rec-LN mice showed increased osteoclastic activity compared to the Virgins but not to the Rec-NN group. On the other hand, maintenance on low protein diet during the recovery period resulted in higher numbers of bone-resorbing cells when compared to control and Rec-NN mice (Fig. 4D, E and F).

**Figure 4.**
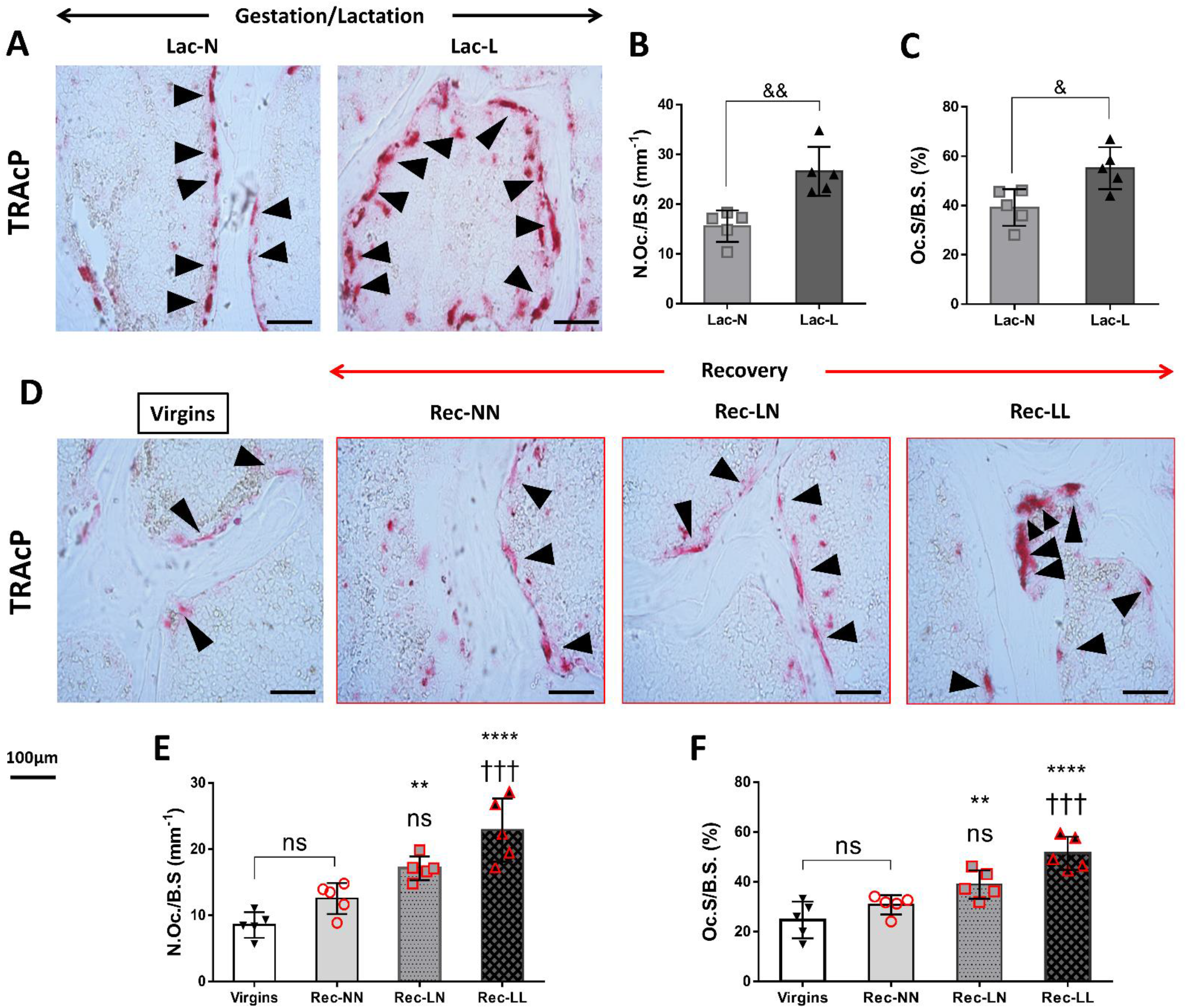
Osteoclastic TRAcP was visualised in Lac-N and Lac-L dams during gestation/lactation (A) and quantified using the number of osteoclasts per bone surface (N.Oc./B.S.) (B) and the percentage of osteoclast surface per bone surface (Oc.S/B.S.) (C) parameters. Similarly, representative images (D) and data comparisons (E and F) are shown for the recovery period against Virgins. Arrows indicate TRAcP-positive osteoclasts attached to the bone surface. All data are presented as mean±SD. ns: not significant; ^&^*p*<0.05, ^&&^*p*<0.01 in Lac-N versus Lac-L comparisons. Asterisks indicate comparisons versus Virgins and crosses versus Rec-NN. \**p*<0.05, \*\**p*<0.01, \*\*\**p*<0.001, \*\*\*\**p*<0.0001.

### Low protein diet reduces *in vitro* bone formation

The next sets of experiments were designed to assess the osteogenic capacity of osteoblasts *in vitro*. Primary osteoblasts were isolated from long-bones of animals of all six groups (n=3/group) and the level of mineralisation using ARS staining was evaluated after 24 days of culture in osteogenic medium. We found that the mineralisation level correlated with the level of protein in the diet that the mice consumed by the end of lactation. Osteoblasts isolated from Lac-N lactating dams formed larger and more numerous bone nodules as compared to the Lac-L group (Fig. 5A and C). Furthermore, mRNA expression levels of *Runx-2*, the master osteogenic transcription factor, as well as *Alp* and *Col1a1*, were all decreased in the Lac-L in comparison with Lac-N (Fig. 5D, G and H), indicating decreased osteoblastic differentiation and activity.

**Figure 5.**
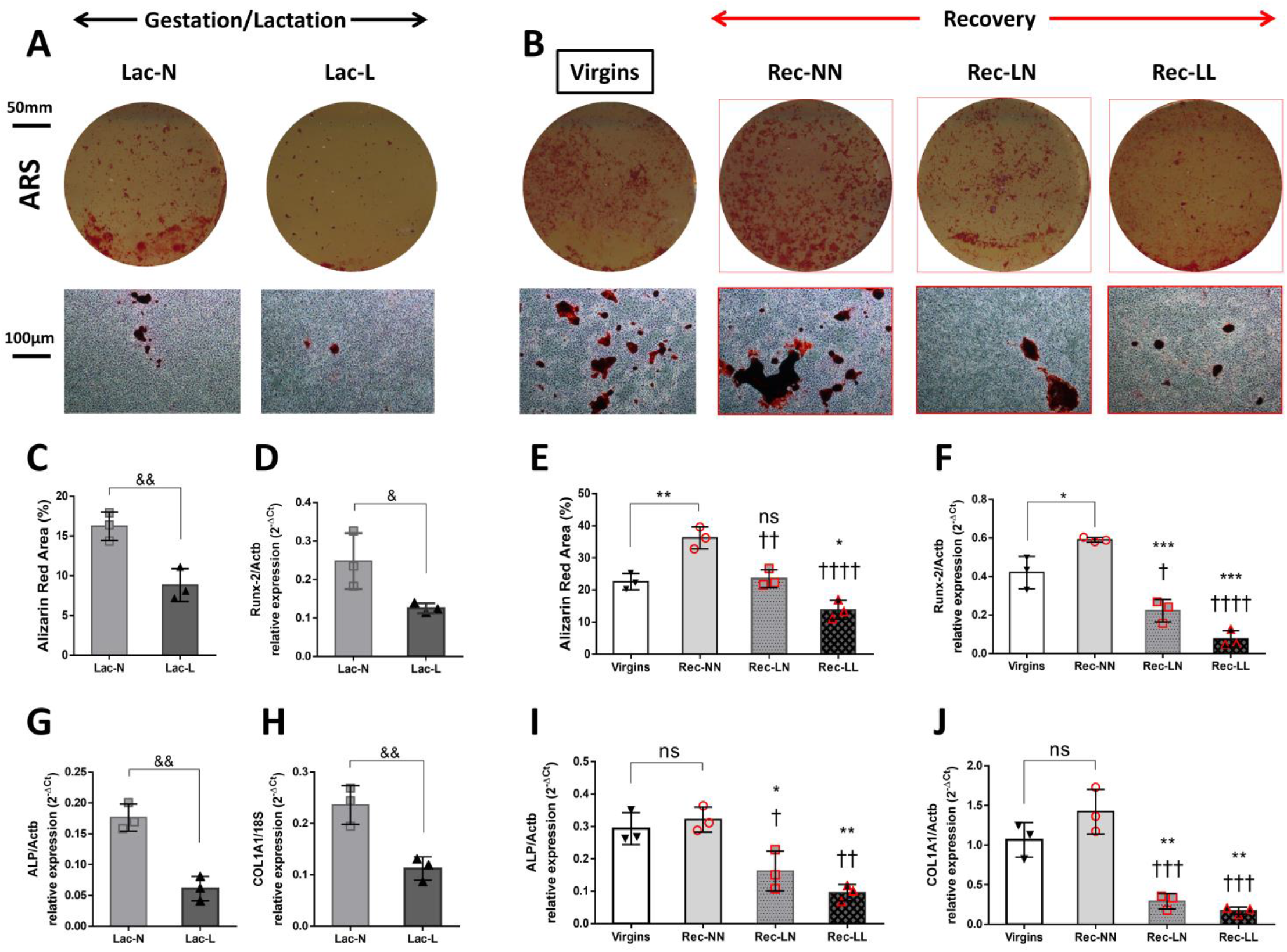
Primary osteoblasts form all the experimental group (n=3/group) were isolated and cultured in osteogenic medium for 4 weeks. Alizarin Res S (ARS) staining (A) was used and quantified (C) for bone mineralisation evaluation and Runx-2 (D), Alp (G) and Col1a1 (H) gene expression was measured by qPCR to assess osteogenic capacity in Lac-N and Lac-L mouse dams during gestation/lactation The corresponding ARS stained bone nodules, ARS quantification and gene expression for the recovery period in Rec-NN, Rec-LN and Rec-LL groups are shown in B, E, F, I and J, respectively, and were compared to the Virgins control. All data are presented as mean±SD. ns: not significant; ^&^*p*<0.05, ^&&^*p*<0.01 in Lac-N versus Lac-L comparisons. Asterisks indicate comparisons versus Virgins and crosses versus Rec-NN. \**p*<0.05, \*\**p*<0.01, \*\*\**p*<0.001, \*\*\*\**p*<0.0001.

In the recovery period, Obs from the Rec-NN group formed a significantly increased number of mineralised bone nodules compared with Virgins and also with Rec-LN and Rec-LL, in agreement with the histological as well as the microCT observations (Fig. 5B and E). Interestingly, mRNA levels of *Alp*, *Col1a1* and *Runx-2*, reflecting osteogenic differentiation and activity, were found increased in the Rec-NN group, and suppressed predominantly in the Rec-LL group (Fig 5F, I and J).

### DNA methylation analysis

To identify potential epigenetic mediators of these effects, we performed targeted pyrosequencing for DNA methylation analysis of the promoters for the *Bmpr1a, Ptch1, Sirt1, Osx* and *Igf1r* genes, while *Runx-2, Alp, Col1a1* were only used as osteogenic markers. The expression patterns of these genes are known to be affected by diet (53–57). They are also considered as crucial molecular players in the major signaling pathways that control osteoblastic differentiation and activity: *Bmpr1a* in BMP pathway, *Ptch1* in hedgehog pathway, *Igf1r* in IGF signaling, while *Osx* is a master regulator of osteoblast differentiation and *Sirt1* links nutritional diet with bone formation (58). All the samples demonstrated negligible DNA methylation, consistent with the promoters being in the active state, with signal ranging within the established noise area of the technology (59). Representative pyrograms are given in Supplementary Figure 2.

### Differential expression of specific miRs may regulate bone recovery delay induced by low protein diet

Based on the *in vitro* results, we hypothesised that the delay of bone recovery in the Rec-LL and, partially, in the Rec-LN group might be caused by differential expression (DE) of bone-related miRs. It has been shown that, among others, some miRs were directly related with bone metabolism. For example, miR-26a regulates osteogenic differentiation of BMSCs and ADSCs by differentially activating Wnt and BMP signaling pathways (60). Furthermore, it has been shown that miR-26a attenuates osteoclastogenesis, actin-ring formation, and bone resorption by suppressing the expression of connective tissue growth factor/CCN family 2 (CTGF/CCN2) (61); miR-34a is down-regulated during osteoclast differentiation (62) and miR-125b controls the osteogenic differentiation of hBMSCs by targeting BMPR1b (63) and also inhibits BMP-4-induced osteoblastic differentiation by regulating cell proliferation in mouse ST2 MSCs, via targeting of the receptor tyrosine kinase Erb2 (64). Therefore, we selected these three miRs, −26a, 34a and 125b, which according to the literature have direct effects on osteoblastic differentiation (60, 62, 63), and performed qPCR to determine their expression levels in Obs isolated from the mouse groups used for the bone recovery period. The levels of miR-26a were found slightly increased in the Rec-NN group as compared to the Virgins and significantly suppressed in the Rec-LN and Rec-LL mice. The endogenous expression of miR-26a is increased during the osteogenic differentiation (65), thus the suppressed levels in Rec-LN and Rec-LL are consistent with decreased osteogenesis. We also found that miR-34a was down-regulated in the Rec-LN and Rec-LL and in-keeping with previous studies reporting that miR-34a-overexpressing transgenic mice exhibit lower bone resorption and higher bone mass through transforming growth factor-β-induced factor 2 (Tgif2) inhibition (62) (62), it is revealed that miR-34a is important for bone regulation. The levels of miR-125b followed the opposite pattern and were elevated in the low bone-forming capacity Obs from the Rec-LN and Rec-LL dams in comparison with the Rec-NN and Virgins groups (Fig. 6A). This was the most important miR that we measured as it directly inhibits Runx-2 (66) and suppresses bone formation by repressing Wnt/β-catenin negative regulators (67) and is directly linked to our findings for decreased expression levels of Runx-2 in the Rec-LL group (Fig. 5).

**Figure 6.**
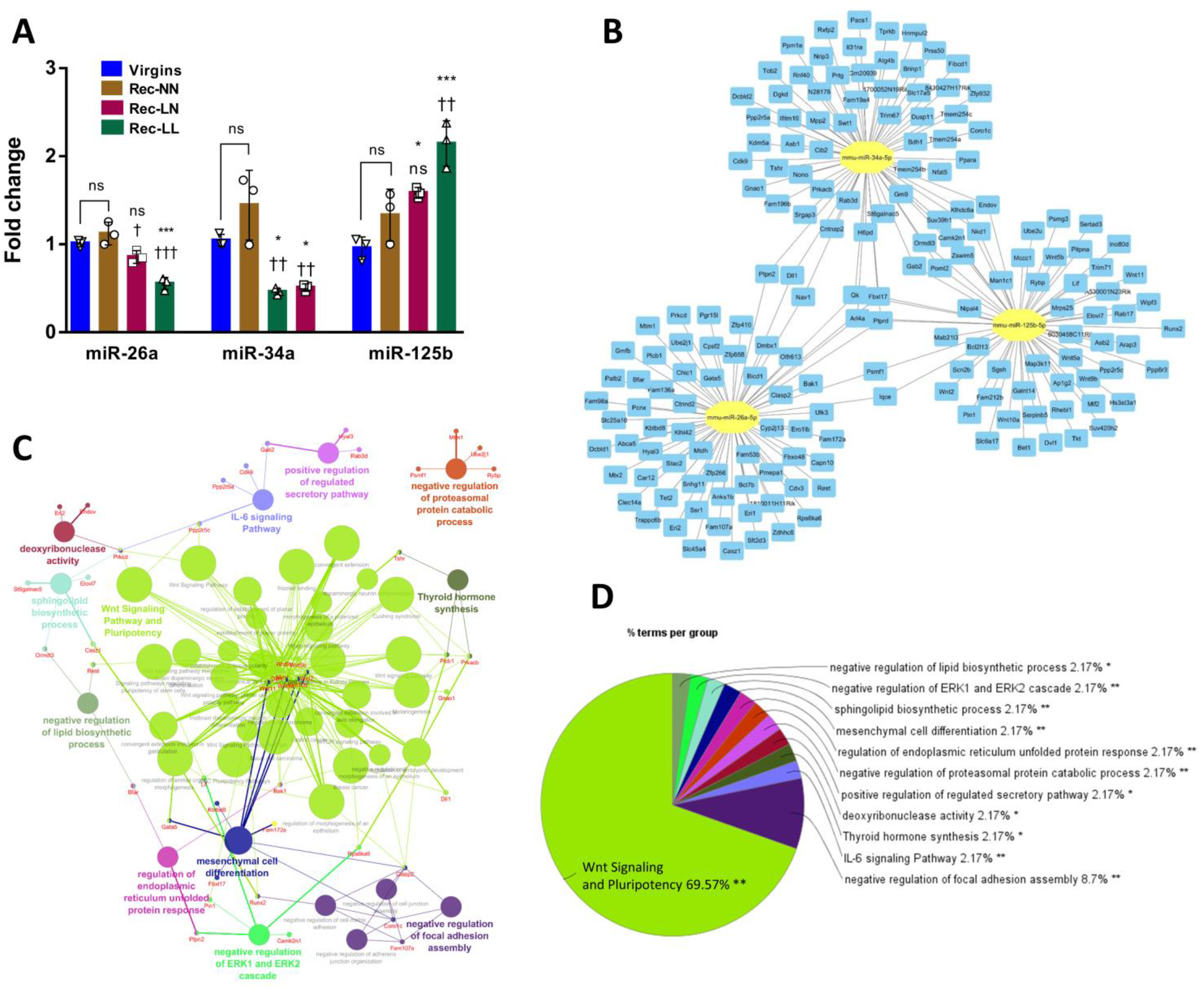
Differential expressions (DE) of miR-26a, 34a and-125b (A) in Rec-NN, Rec-LN and Rec-LL groups are shown for the recovery period and were compared to the Virgins control (n=3/group). Using bioinformatics, the predicted target genes were mapped (B) and biomolecular interactions were designed by gene ontology (GO) and KEGG pathways analyses (C). Wnt signaling seems to be the dominant pathway affected by the selected miRs among the statistically significant regulatory mechanisms (D). All data are presented as mean±SD. ns: not significant. Asterisks indicate comparisons versus Virgins and crosses versus Rec-NN. \**p*<0.05, \*\**p*<0.01, \*\*\**p*<0.001.

Using the DE of these specific miRs, bioinformatic analyses revealed a total of 174 genes predicted as potential targets, while 18 genes were common (Fig. 6B, enlargement in Supplementary Figure 3). Statistically significant biomolecular interactions, using constructed networks of GO and KEGG pathways, showed that, among others, some important bone regulatory mechanisms and biological processes can be affected, namely the Wnt and IL-6 signaling pathways, thyroid hormone synthesis, mesenchymal cell differentiation and negative regulation of ERK1 and ERK2 cascades (Fig. 6C, enlargement in Supplementary Figure 4). Finally, it was found that the vast majority of the implicated gene targets (69.57%) were involved in Wnt signaling and pluripotency (Fig. 6D).

## DISCUSSION

In this study we report that protein under-nutrition during gestation/lactation and recovery periods reproduction exerts detrimental effects on the skeletal system of mouse dams and decreases the rate of bone accretion in the post-weaning recovery period. We also examined the effects of low protein intake during gestation and lactation on skeletal muscle integrity as well as on NMJ morphology and importantly report for the first time in the literature that there were no differences when compared with normal protein consumption.

Our first aim was to explore if a low protein diet during gestation and lactation can affect skeletal muscles. Lactation is linked to significant changes in maternal metabolism and subsequent adaptations, such as decreased adaptive thermogenesis, are necessary due to high-energy demands for sufficient milk production (68, 69). Liver, skeletal muscle and white adipose tissue are the main metabolic tissues in mammals (70), and thus their metabolic rates are modulated during lactation. Skeletal muscle protein mobilisation acts as an adaptive response to meet these energy requirements and is also finely regulated in proportion to the dietary protein intake. In our cohort of mouse dams, no significant change in the TA skeletal muscle fibre CSA was observed, possibly due to activation of other metabolic pathways aiming to fulfil energy demands since both experimental diets (normal and low) were isocaloric. Maternal protein loss during lactation is decreased when total body protein mass (10, 11) exceeds certain levels to prevent exhaustion (71). Similarly, no differences were observed in NMJ morphology and structure between Virgins, Lac-N and Lac-L groups, supporting our findings that low protein intake during gestation and lactation does not dramatically alter skeletal muscles as these two systems function integrally.

Our second aim was to determine changes in the skeleton of mouse dams due to low protein diet during gestation and lactation. Pregnancy and lactation are very challenging periods for the maternal skeleton due to significant adjustments in calcium homeostasis. Our study showed that lactation resulted in a significant decrease of bone volume and deterioration of bone micro-architecture in both trabecular and cortical bone. It is known that lactation causes demineralization of the maternal skeleton which leads to bone mass reduction (72). BV/TV was significantly reduced in the proximal tibia and L5 vertebral body, and this was accompanied by thinning of the trabeculae at both skeletal sites. Cortical bone thickness at the tibial mid-diaphysis was also considerably decreased by lactation. These changes were exacerbated in the Lac-L mouse dams indicating that low protein consumption leads to more profound skeletal deterioration than normal protein consumption during the reproductive period. Although some studies have reported that the extend of lactation-induced bone loss is different according to the anatomical site of the skeleton and is, particularly, higher in the spine than in the long-bones (19), our model did not identify any differences showing a universal effect on both sites. An important finding was that all microCT structural parameters were worsened in the Lac-L group as compared to Lac-N, and the changes in SMI and Tb.N show that low protein diet affects not only the overall bone mass but also leads to micro-architectural changes. The increased SMI shows that the shape of trabeculae has altered from plate-like to a more rod-like structure, similar to other studies (73). However, these studies have significant differences from the present study, i.e. mouse strain and larger litter size. Histomorphometric analyses of animal models have shown that bone turnover is increased during pregnancy (74) and these high rates of bone formation and resorption are maintained throughout lactation. Our results indicate that mouse dams fed a normal diet during gestation and lactation, Lac-N, follow this trend but, on the contrary, the Lac-L mice show suppressed numbers of osteoblasts, and very high levels of osteoclastic bone resorption. Thus, the overall result is a significant bone loss as compared to the Lac-N group, and this is supported by the results of the microCT analysis. Our *in vitro* findings support our hypothesis that enhanced bone loss in the Lac-L mice was caused, at least partly, by the substantially reduced osteogenic capacity of the osteoblasts that was coupled with decreased expression of osteogenic genes in comparison with the Lac-N group. Dietary protein has beneficial effects on bone health (75), but very little is known regarding lactation. The mechanisms that regulate lactation-related skeletal loss have been extensively studied, suggesting an interplay between increased osteocytic osteolysis, coordination by the brain-breast-bone circuit through pituitary-derived prolactin and oxytocin in response to suckling which leads to the release of parathyroid hormone-related protein (PTHrP) and subsequent proliferation and activation of osteoclasts (72). However, the effects of protein intake on this molecular network are largely unknown and this needs further investigation.

Our next goal was to study the effects of low protein diet on the bone recovery period after weaning. Regardless of the accelerated bone loss that occurs during lactation, bonemass recovers promptly after weaning and termination of milk production. The recovery process is characterized by a sudden cessation of bone resorption and a remarkably elevated rate of bone formation with rapid re-mineralisation of new bone matrix (15, 76). Normally, the recovery period lasts approximately 6 months in humans and 4 weeks in mice and mineral content reaches the baseline values before pregnancy by gaining 2% to 3% mineral apposition per month in humans and 10% to 20% per week in mice (1, 77). However, some studies suggest that pharmaceutical approaches such as zoledronate (73) or osteoprotegerin (78) administration can prevent maternal lactation-induced bone loss and improve bone recovery, but concerns may arise for neonatal fetal growth and health. We found that after 28 days from the end of lactation only the dams that were on the normal protein diet throughout the entire experimental period (Rec-NN) achieved full recovery with microCT bone parameters that did not differ from the nulliparous control at any of the skeletal anatomical sites. Rec-LN mice had lower bone mass compared to the Rec-NN group which shows that switching from low to normal protein diet after weaning leads to only partial bone recovery. The difference between Rec-LN and Rec-LL was of great interest, showing that maintaining mice on a low protein diet leads to an extensive delay in skeletal recovery. Interestingly, the Rec-LL group did not differ from the Lac-L during lactation, showing signs of persistent failure to recuperate lactation-induced bone reduction.

Our results indicate that a low protein diet has deleterious effects on osteoblasts and enhances osteoclastic bone resorption. Elefteriou *et al.* (79) have shown that activating transcription factor 4 (ATF4), a transcription factor that enhances amino-acid uptake and collagen synthesis in osteoblasts, is a crucial player highlighting the significance of protein intake in bone formation. ATF4^−/−^ mice showed a deformed skeletal phenotype that was rescued by high protein diet intake through an increase of collagen type I synthesis and osteocalcin expression by differentiated osteoblasts due to higher amino-acid uptake. It has also been reported that protein malnutrition stimulates bone marrow mesenchymal stem cells differentiation to adipocytes rather than osteoblasts (80) and attenuates the bone anabolic response to PTH in female rats (81, 82). Therefore, we examined the behaviour of isolated primary Obs from the animals used for the recovery period experiments. The effect of low protein diet on osteoblasts was verified by our *in vitro* bone formation assay and gene expression of osteogenic markers. These results were unexpected since all cell cultures were managed under the same experimental conditions, i.e. equal serum concentration and all in osteogenic medium. We assumed that when isolated Obs return to normal protein levels *in vitro*, they would have similar osteogenic behaviour. In contrast with our expectations, osteoblastic cells retained the *in vivo* bone-forming capacity. These striking results led us to the novel conclusion that the nutritional protein level is able to leave a molecular signature in these cells.

We, therefore, hypothesised that the observed osteogenic activity during the recovery period is potentially regulated by epigenetic mechanisms or differential expression of bone-related miRs. We selected four key genes that are master regulators of osteoblastic differentiation and activation (*Bmpr1a, Ptch1, Osx* and *Igf1r*) (83) and *Sirt1,* a NAD-dependent deacetylase, which promotes osteogenic differentiation (84) and its expression is affected by dietary protein (57). Specific-site DNA pyrosequencing analysis revealed no differences in methylation patterns of selected genomic loci in the promoter region of osteoblastic regulatory genes. However, other CpG rich regions may be affected and this possibility needs further study. On the other hand, we found differential expression of miRs −26a, −34a and −125b which have been shown to regulate bone-related genes. MiR-125b regulates the osteogenic differentiation of human MSCs by targeting BMPR1b (63) while miR-26a reverses the bone regeneration deficit of MSCs (85), and miR-34a inhibits osteoclastogenesis (62). During the bone recovery period, the expression of these miRs was altered favouring suppression of bone formation in combination with elevated resorption that was profound in the Rec-LL mice. This effect can be driven by dietary protein content as the Rec-LN dams showed partially recovery. Based on these observations, we performed a bioinformatic analysis aiming to examine which genes could be potential targets of the specific miRs, and which revealed regulatory pathways involved in important bone homeostatic mechanisms. The main pathways affected appear to be the Wnt and IL-6 signaling pathways, and these have profound effects on osteoblasts and osteoclasts respectively (86, 87).

In conclusion, here we report that low protein intake during the reproduction period does not affect the skeletal muscles and associated NMJs in mouse dams. However, protein under-nutrition increases lactation-induced bone loss, and maintenance on low protein diet during the recovery period delays bone restoration. Importantly, isolated osteoblasts show similar osteogenic behaviour to the *in vivo* findings, suggesting a possible epigenetic mechanism. This dietary protein-dependent effect on bone metabolism might be controlled by changes in the expression of specific miRs. Further studies are required to identify the mechanism(s) underlying these effects of low protein diet. A full understanding of the mechanisms would be expected to lead to improved nutritional guidelines during reproduction and could identify new targets for the treatment of musculoskeletal disorders.

## ACKNOWLEDGEMENTS

This work was funded by the BBSRC (BB/P008429/1) to A.V. and K.G-W., and departmental support from the Institute of Life Course and Medical Sciences, University of Liverpool to I.K. We also thank the Biomedical Services Unit at the University of Liverpool. S.E.O. is a member of the MRC Metabolic Diseases Unit (MC_UU_00014/4).

## AUTHOR CONTRIBUTIONS

I.K. and A.V. designed the experiments with input from K.G-W., R.J.H., T.L., A.S. and S.E.O.; I.K., M.A. and M.S. performed the experiments, acquired and analysed the data; I.K. and A.V. wrote the manuscript, which was critically revised and approved by all co-authors.

---

**Supplementary Figure 1.**
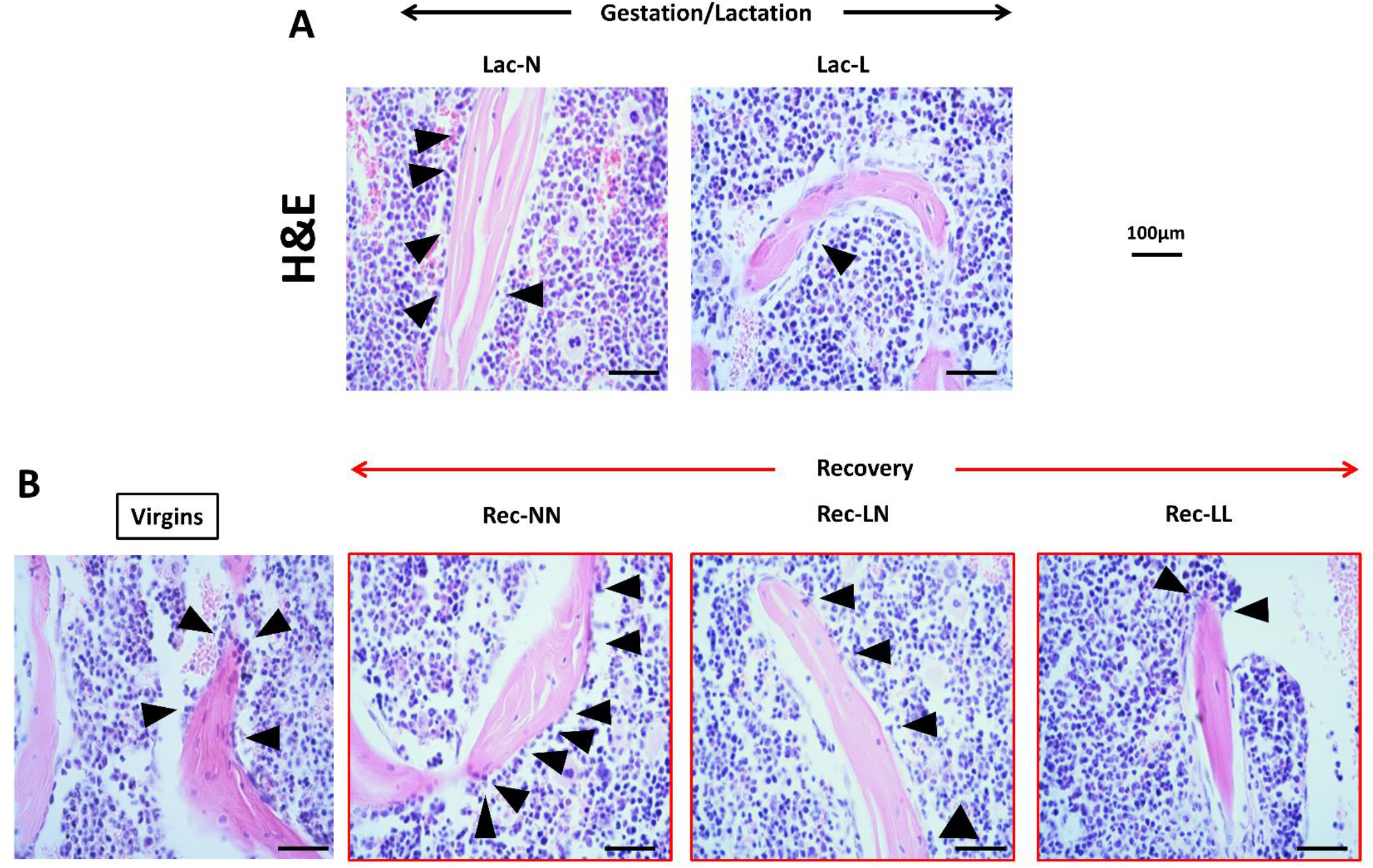
Representative H&E stained L5 vertebral body sections Lac-N and Lac-L mouse dams during gestation/lactation (A) and for the recovery period in Rec-NN, Rec-LN and Rec-LL (B) groups are shown in comparison to the Virgins control. Arrows indicate osteoblasts attached to the bone surface.

**Supplementary Figure 2.**
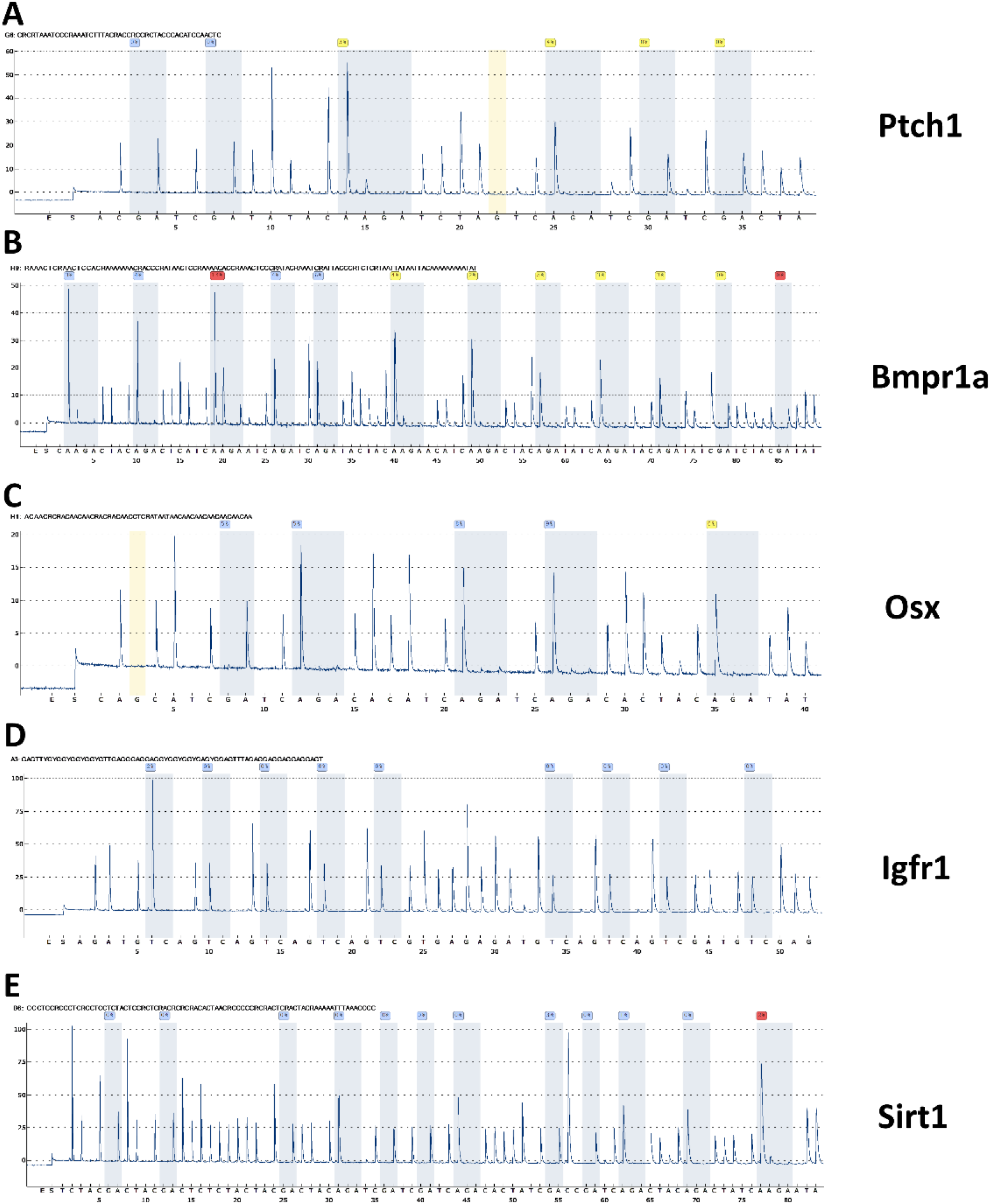
Representative pyrograms for *Ptch1* (A), *Bmpr1a* (B), *Osx* (C), *Igfr1* (D) and *Sirt1* (E) from pyrosequencing based DNA methylation analysis. X axis demonstrates the dispensation order for each assay. Blue lanes indicate the interrogated CpG sites. The software calculated DNA methylation of each site as the percentage of C/[C+T] (or G/[A+G] in reverse sequencing) incorporation. Y axis represents the light signal produced by a coupled reaction following nucleotide incorporation.

**Supplementary Figure 3.**
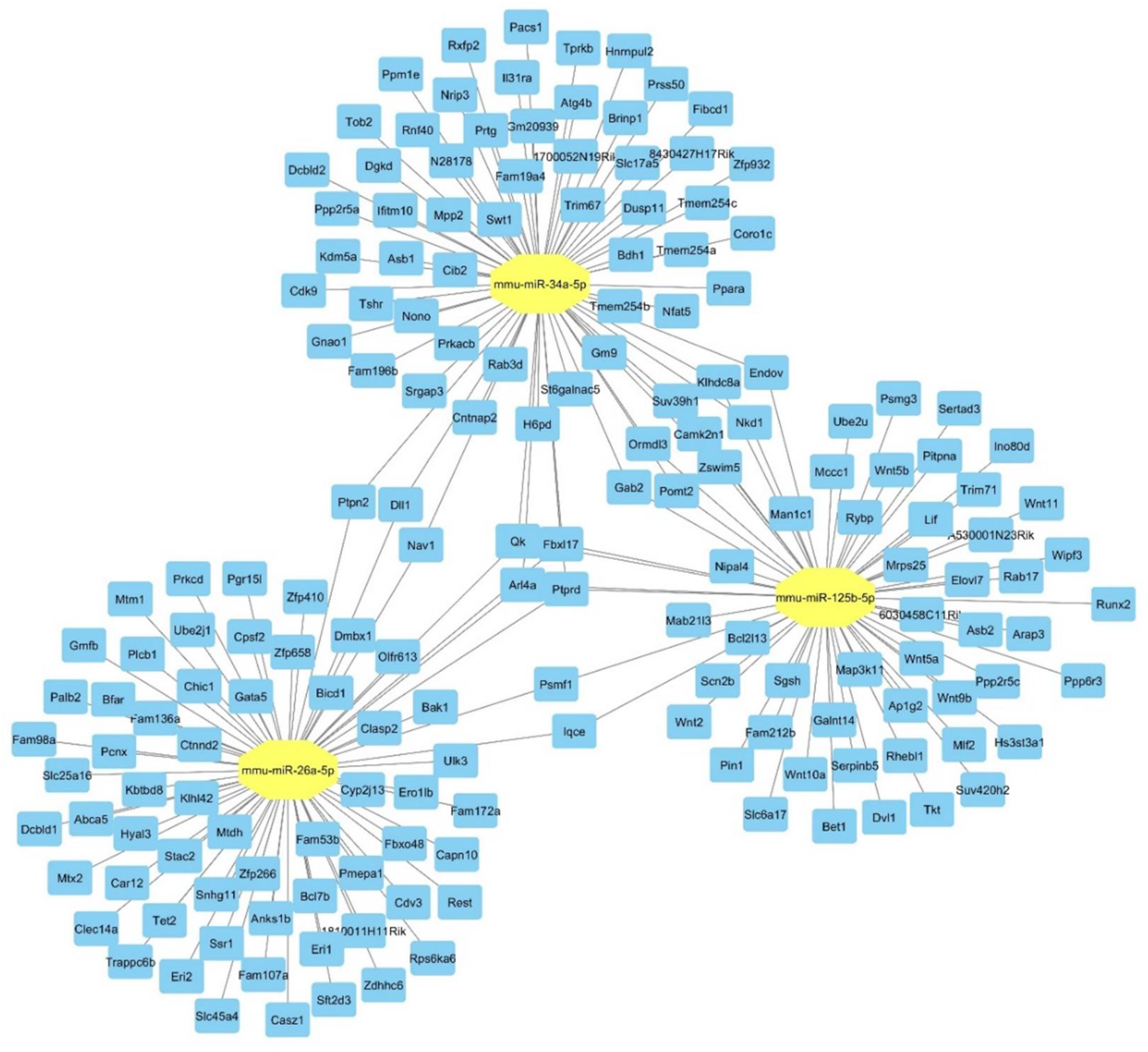
Enlargement of Figure 6B for better visualisation of predicted gene targets of miR-26a, −34a and −125b.

**Supplementary Figure 4.**
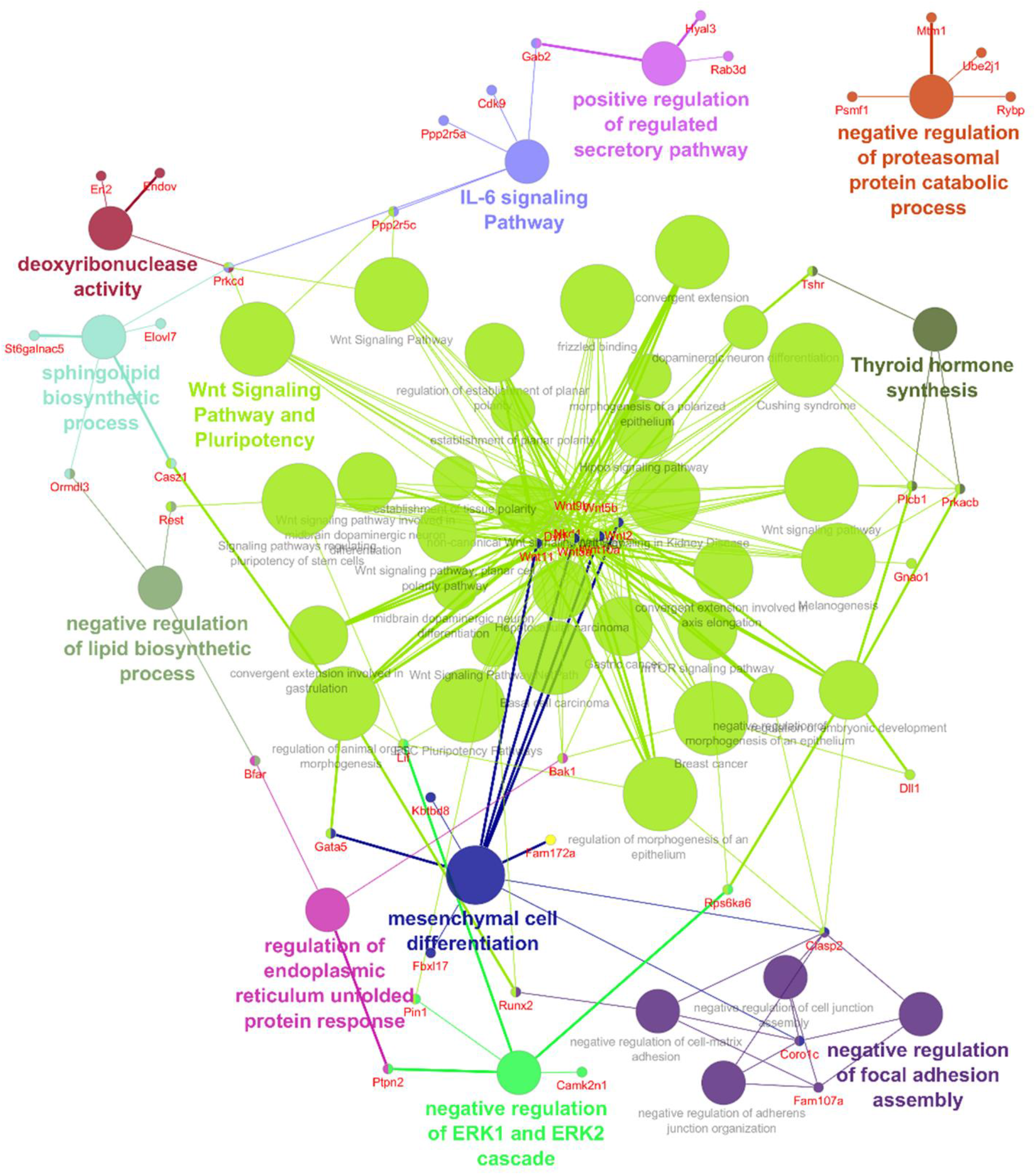
Enlargement of Figure 6C for better visualisation of the molecular pathway interactions of the predicted gene targets of miR-26a, −34a and −125b.

**Supplementary Table 1:**
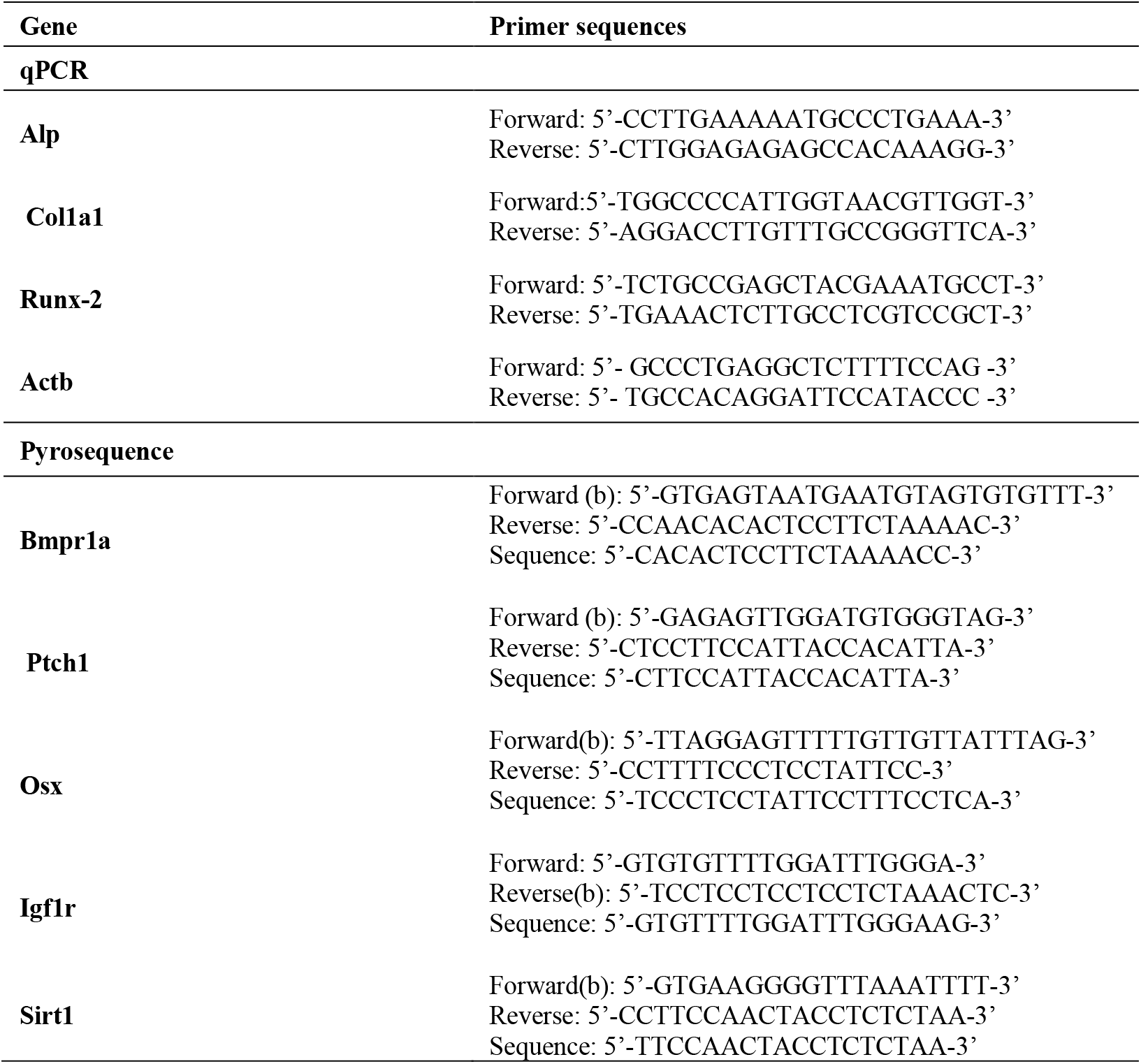
Sequences of the primers used for qPCR and pyrosequencing; b: biotinylated primer.

**Supplementary Table 2:** Trabecular bone morphometry of proximal tibiae and L5 vertebral body analysed by microCT. TV: tissue volume; BV/TV: bone volume to tissue volume ratio; Tb.Th: trabecular thickness; Tb.Sp: trabecular separation; Tb.N: trabecular number; SMI: structure model index, Tb.Pf: trabecular pattern factor. All data shown are means±SD. ns: not significant; ^&^*p*<0.05, ^&&^*p*<0.01, ^&&&^*p*<0.001, ^&&&&^*p*<0.0001 in Lac-N versus Lac-L comparisons. Asterisks indicate comparisons versus Virgins and crosses versus Rec-NN.

| Gestation/Lactation |  |  | Virgins<br>(n=8) | Recovery |  |  |
| --- | --- | --- | --- | --- | --- | --- |
| Tibia | Lac-N<br>(n=8) | Lac-L<br>(n=8) |  | Rec-NN<br>(n=5) | Rec-LN<br>(n=5) | Rec-LL<br>(n=5) |
| TV (mm<sup>3</sup>) | 3.33±0.29 | 3.19±0.47 | 3.18±0.21 | 3.39±0.49 | 3.44±0.14 | 3.28±0.33 |
| Tb.Th (µm) | 49.17±3.51 | 43.83±1.94<sup>&</sup> | 57.10±3.09 | 59.28±2.71 | 55.82±4.02 | 47.25±1.73<sup>****, ††††</sup> |
| Tb.Sp (µm) | 419.88±25.06 | 529.92±86.83<sup>&&</sup> | 322.50±22.13 | 314.02±24.77 | 359.35±29.70<sup>**, ††</sup> | 516.05±59.29<sup>****, †††</sup> |
| Tb.N (mm<sup>-1</sup>) | 1.00±0.13 | 0.49±0.15<sup>&&&</sup> | 3.11±0.12 | 3.20±0.37 | 1.42±0.09<sup>***, ††††</sup> | 0.70±0.13<sup>****, ††††</sup> |
| SMI | 2.51±0.14 | 2.84±0.13<sup>&&</sup> | 2.15±0.13 | 2.17±0.20 | 2.41±0.14<sup>*, †</sup> | 2.74±0.11<sup>****, ††††</sup> |
| Tb.Pf (mm<sup>-1</sup>) | 31.32±2.43 | 35.21±1.47<sup>&&</sup> | 27.58±2.83 | 25.30±1.87 | 31.96±1.43<sup>**, ††</sup> | 35.05±2.01<sup>**, †††</sup> |
| **L5** |  |  |  |  |  |  |
| TV (mm<sup>3</sup>) | 3.51±0.29 | 3.55±0.18 | 3.53±0.12 | 3.37±0.20 | 3.51±0.12 | 3.32±0.13 |
| Tb.Th (µm) | 54.77±5.15 | 47.11±4.85<sup>&</sup> | 62.67±3.16<sup>*, ††</sup> | 60.22±2.37 | 54.65±4.38<sup>*, ns</sup> | 49.34±4.09<sup>***, ††</sup> |
| Tb.Sp (µm) | 317.7±29.31 | 373.5±16.64<sup>&&</sup> | 265.3±30.95<sup>*, ††</sup> | 256.3±25.43 | 311.5±19.00<sup>**, ††</sup> | 362.1±35.81<sup>****, ††††</sup> |
| Tb.N (mm<sup>-1</sup>) | 3.26±0.21 | 2.98±0.32<sup>&</sup> | 4.45±0.34 | 4.33±0.41 | 3.91±0.22<sup>*, †</sup> | 3.33±0.45<sup>****, †††</sup> |
| SMI | 1.91±0.13 | 2.34±0.09<sup>&&</sup> | 1.40±0.05 | 1.44±0.07 | 1.72±0.09<sup>*, †</sup> | 2.37±0.15<sup>***, †††</sup> |
| Tb.Pf (mm<sup>-1</sup>) | 33.54±1.65 | 38±1.52<sup>&&</sup> | 22.85±1.58 | 23.87±2.22 | 29±2.16<sup>*, ns</sup> | 37.68±1.98<sup>****, †††</sup> |

## REFERENCES

1. Kovacs, C. S., and Kronenberg, H. M. (1997) Maternal-fetal calcium and bone metabolism during pregnancy, puerperium, and lactation. Endocr Rev 18, 832–872

2. Kovacs, C. S. (2011) Calcium and bone metabolism disorders during pregnancy and lactation. Endocrinol Metab Clin North Am 40, 795–826

3. Kovacs, C. S., and Fuleihan Gel, H. (2006) Calcium and bone disorders during pregnancy and lactation. Endocrinol Metab Clin North Am 35, 21–51, v

4. Kovacs, C. S. (2005) Calcium and bone metabolism during pregnancy and lactation. J Mammary Gland Biol Neoplasia 10, 105–118

5. Olausson, H., Goldberg, G. R., Laskey, M. A., Schoenmakers, I., Jarjou, L. M., and Prentice, A. (2012) Calcium economy in human pregnancy and lactation. Nutr Res Rev 25, 40–67

6. Prentice, A. (2003) Micronutrients and the bone mineral content of the mother, fetus and newborn. J Nutr 133, 1693S–1699S

7. Sowers, M., Corton, G., Shapiro, B., Jannausch, M. L., Crutchfield, M., Smith, M. L., Randolph, J. F., and Hollis, B. (1993) Changes in bone density with lactation. JAMA 269, 3130–3135

8. Kent, G. N., Price, R. I., Gutteridge, D. H., Smith, M., Allen, J. R., Bhagat, C. I., Barnes, M. P., Hickling, C. J., Retallack, R. W., Wilson, S. G., and et al. (1990) Human lactation: forearm trabecular bone loss, increased bone turnover, and renal conservation of calcium and inorganic phosphate with recovery of bone mass following weaning. J Bone Miner Res 5, 361–369

9. Zeni, S. N., Di Gregorio, S., and Mautalen, C. (1999) Bone mass changes during pregnancy and lactation in the rat. Bone 25, 681–685

10. Ardeshirpour, L., Dann, P., Adams, D. J., Nelson, T., VanHouten, J., Horowitz, M. C., and Wysolmerski, J. J. (2007) Weaning triggers a decrease in receptor activator of nuclear factor-kappaB ligand expression, widespread osteoclast apoptosis, and rapid recovery of bone mass after lactation in mice. Endocrinology 148, 3875–3886

11. Bowman, B. M., Siska, C. C., and Miller, S. C. (2002) Greatly increased cancellous bone formation with rapid improvements in bone structure in the rat maternal skeleton after lactation. J Bone Miner Res 17, 1954–1960

12. Vajda, E. G., Bowman, B. M., and Miller, S. C. (2001) Cancellous and cortical bone mechanical properties and tissue dynamics during pregnancy, lactation, and postlactation in the rat. Biol Reprod 65, 689–695

13. Paton, L. M., Alexander, J. L., Nowson, C. A., Margerison, C., Frame, M. G., Kaymakci, B., and Wark, J. D. (2003) Pregnancy and lactation have no long-term deleterious effect on measures of bone mineral in healthy women: a twin study. Am J Clin Nutr 77, 707–714

14. Melton, L. J., 3rd, Bryant, S. C., Wahner, H. W., O’Fallon, W. M., Malkasian, G. D., Judd, H. L., and Riggs, B. L. (1993) Influence of breastfeeding and other reproductive factors on bone mass later in life. Osteoporos Int 3, 76–83

15. Lissner, L., Bengtsson, C., and Hansson, T. (1991) Bone mineral content in relation to lactation history in pre- and postmenopausal women. Calcif Tissue Int 48, 319–325

16. Hopkinson, J. M., Butte, N. F., Ellis, K., and Smith, E. O. (2000) Lactation delays postpartum bone mineral accretion and temporarily alters its regional distribution in women. J Nutr 130, 777–783

17. Karlsson, C., Obrant, K. J., and Karlsson, M. (2001) Pregnancy and lactation confer reversible bone loss in humans. Osteoporos Int 12, 828–834

18. Pearson, D., Kaur, M., San, P., Lawson, N., Baker, P., and Hosking, D. (2004) Recovery of pregnancy mediated bone loss during lactation. Bone 34, 570–578

19. Kirby, B. J., Ardeshirpour, L., Woodrow, J. P., Wysolmerski, J. J., Sims, N. A., Karaplis, A. C., and Kovacs, C. S. (2011) Skeletal recovery after weaning does not require PTHrP. J Bone Miner Res 26, 1242–1251

20. Clowes, E. J., Aherne, F. X., and Baracos, V. E. (2005) Skeletal muscle protein mobilization during the progression of lactation. Am J Physiol Endocrinol Metab 288, E564–572

21. Bell, A. W., Burhans, W. S., and Overton, T. R. (2000) Protein nutrition in late pregnancy, maternal protein reserves and lactation performance in dairy cows. Proc Nutr Soc 59, 119–126

22. Kuhla, B., Nurnberg, G., Albrecht, D., Gors, S., Hammon, H. M., and Metges, C. C. (2011) Involvement of skeletal muscle protein, glycogen, and fat metabolism in the adaptation on early lactation of dairy cows. J Proteome Res 10, 4252–4262

23. Thomas, M., and Weisman, S. M. (2006) Calcium supplementation during pregnancy and lactation: effects on the mother and the fetus. Am J Obstet Gynecol 194, 937–945

24. Grieger, J. A., and Clifton, V. L. (2014) A review of the impact of dietary intakes in human pregnancy on infant birthweight. Nutrients 7, 153–178

25. Peterson, C. A., Schnell, J. D., Kubas, K. L., and Rottinghaus, G. E. (2009) Effects of soy isoflavone consumption on bone structure and milk mineral concentration in a rat model of lactation-associated bone loss. Eur J Nutr 48, 84–91

26. Bueno-Vargas, P., Manzano, M., Diaz-Castro, J., Lopez-Aliaga, I., Rueda, R., and Lopez-Pedrosa, J. M. (2016) Maternal Dietary Supplementation with Oligofructose-Enriched Inulin in Gestating/Lactating Rats Preserves Maternal Bone and Improves Bone Microarchitecture in Their Offspring. PLoS One 11, e0154120

27. Nishikawa, K., Iwamoto, Y., Kobayashi, Y., Katsuoka, F., Kawaguchi, S., Tsujita, T., Nakamura, T., Kato, S., Yamamoto, M., Takayanagi, H., and Ishii, M. (2015) DNA methyltransferase 3a regulates osteoclast differentiation by coupling to an S-adenosylmethionine-producing metabolic pathway. Nat Med 21, 281–287

28. Delgado-Calle, J., and Riancho, J. A. (2012) The role of DNA methylation in common skeletal disorders. Biology (Basel) 1, 698–713

29. Delgado-Calle, J., Sanudo, C., Bolado, A., Fernandez, A. F., Arozamena, J., Pascual-Carra, M. A., Rodriguez-Rey, J. C., Fraga, M. F., Bonewald, L., and Riancho, J. A. (2012) DNA methylation contributes to the regulation of sclerostin expression in human osteocytes. J Bone Miner Res 27, 926–937

30. Delgado-Calle, J., Sanudo, C., Fernandez, A. F., Garcia-Renedo, R., Fraga, M. F., and Riancho, J. A. (2012) Role of DNA methylation in the regulation of the RANKL-OPG system in human bone. Epigenetics 7, 83–91

31. Ghayor, C., and Weber, F. E. (2016) Epigenetic Regulation of Bone Remodeling and Its Impacts in Osteoporosis. Int J Mol Sci 17

32. Zhang, Y., Xie, R. L., Croce, C. M., Stein, J. L., Lian, J. B., van Wijnen, A. J., and Stein, G. S. (2011) A program of microRNAs controls osteogenic lineage progression by targeting transcription factor Runx2. Proc Natl Acad Sci U S A 108, 9863–9868

33. Li, Z., Hassan, M. Q., Volinia, S., van Wijnen, A. J., Stein, J. L., Croce, C. M., Lian, J. B., and Stein, G. S. (2008) A microRNA signature for a BMP2-induced osteoblast lineage commitment program. Proc Natl Acad Sci U S A 105, 13906–13911

34. Huang, J., Zhao, L., Xing, L., and Chen, D. (2010) MicroRNA-204 regulates Runx2 protein expression and mesenchymal progenitor cell differentiation. Stem Cells 28, 357–364

35. Vimalraj, S., Partridge, N. C., and Selvamurugan, N. (2014) A positive role of microRNA-15b on regulation of osteoblast differentiation. J Cell Physiol 229, 1236–1244

36. Hu, R., Liu, W., Li, H., Yang, L., Chen, C., Xia, Z. Y., Guo, L. J., Xie, H., Zhou, H. D., Wu, X. P., and Luo, X. H. (2011) A Runx2/miR-3960/miR-2861 regulatory feedback loop during mouse osteoblast differentiation. J Biol Chem 286, 12328–12339

37. Liu, P., Baumgart, M., Groth, M., Wittmann, J., Jack, H. M., Platzer, M., Tuckermann, J. P., and Baschant, U. (2016) Dicer ablation in osteoblasts by Runx2 driven cre-loxP recombination affects bone integrity, but not glucocorticoid-induced suppression of bone formation. Sci Rep 6, 32112

38. Bendre, A., Moritz, N., Vaananen, V., and Maatta, J. A. (2018) Dicer1 ablation in osterix positive bone forming cells affects cortical bone homeostasis. Bone 106, 139–147

39. Glatt, V., Canalis, E., Stadmeyer, L., and Bouxsein, M. L. (2007) Age-related changes in trabecular architecture differ in female and male C57BL/6J mice. J Bone Miner Res 22, 1197–1207

40. van ‘t Hof, R. J., Rose, L., Bassonga, E., and Daroszewska, A. (2017) Open source software for semi-automated histomorphometry of bone resorption and formation parameters. Bone 99, 69–79

41. Dempster, D. W., Compston, J. E., Drezner, M. K., Glorieux, F. H., Kanis, J. A., Malluche, H., Meunier, P. J., Ott, S. M., Recker, R. R., and Parfitt, A. M. (2013) Standardized nomenclature, symbols, and units for bone histomorphometry: a 2012 update of the report of the ASBMR Histomorphometry Nomenclature Committee. J Bone Miner Res 28, 2–17

42. Kanakis, I., Liu, K., Poulet, B., Javaheri, B., van ‘t Hof, R. J., Pitsillides, A. A., and Bou-Gharios, G. (2019) Targeted Inhibition of Aggrecanases Prevents Articular Cartilage Degradation and Augments Bone Mass in the STR/Ort Mouse Model of Spontaneous Osteoarthritis. Arthritis Rheumatol 71, 571–582

43. van ‘t Hof, R. J. (2012) Analysis of bone architecture in rodents using microcomputed tomography. Methods Mol Biol 816, 461–476

44. Taylor, S. E., Shah, M., and Orriss, I. R. (2014) Generation of rodent and human osteoblasts. Bonekey Rep 3, 585

45. Stephens, A. S., Stephens, S. R., and Morrison, N. A. (2011) Internal control genes for quantitative RT-PCR expression analysis in mouse osteoblasts, osteoclasts and macrophages. BMC Res Notes 4, 410

46. Livak, K. J., and Schmittgen, T. D. (2001) Analysis of relative gene expression data using real-time quantitative PCR and the 2(-Delta Delta C(T)) Method. Methods 25, 402–408

47. Dweep, H., and Gretz, N. (2015) miRWalk2.0: a comprehensive atlas of microRNA-target interactions. Nat Methods 12, 697

48. Agarwal, V., Bell, G. W., Nam, J. W., and Bartel, D. P. (2015) Predicting effective microRNA target sites in mammalian mRNAs. Elife 4

49. Wong, N., and Wang, X. (2015) miRDB: an online resource for microRNA target prediction and functional annotations. Nucleic Acids Res 43, D146–152

50. Hsu, S. D., Lin, F. M., Wu, W. Y., Liang, C., Huang, W. C., Chan, W. L., Tsai, W. T., Chen, G. Z., Lee, C. J., Chiu, C. M., Chien, C. H., Wu, M. C., Huang, C. Y., Tsou, A. P., and Huang, H. D. (2011) miRTarBase: a database curates experimentally validated microRNA-target interactions. Nucleic Acids Res 39, D163–169

51. Shannon, P., Markiel, A., Ozier, O., Baliga, N. S., Wang, J. T., Ramage, D., Amin, N., Schwikowski, B., and Ideker, T. (2003) Cytoscape: a software environment for integrated models of biomolecular interaction networks. Genome Res 13, 2498–2504

52. Kanehisa, M., and Goto, S. (2000) KEGG: kyoto encyclopedia of genes and genomes. Nucleic Acids Res 28, 27–30

53. Schulz, T. J., Graja, A., Huang, T. L., Xue, R., An, D., Poehle-Kronawitter, S., Lynes, M. D., Tolkachov, A., O’Sullivan, L. E., Hirshman, M. F., Schupp, M., Goodyear, L. J., Mishina, Y., and Tseng, Y. H. (2016) Loss of BMP receptor type 1A in murine adipose tissue attenuates age-related onset of insulin resistance. Diabetologia 59, 1769–1777

54. Jalabert, A., Vial, G., Guay, C., Wiklander, O. P., Nordin, J. Z., Aswad, H., Forterre, A., Meugnier, E., Pesenti, S., Regazzi, R., Danty-Berger, E., Ducreux, S., Vidal, H., El-Andaloussi, S., Rieusset, J., and Rome, S. (2016) Exosome-like vesicles released from lipid-induced insulin-resistant muscles modulate gene expression and proliferation of beta recipient cells in mice. Diabetologia 59, 1049–1058

55. Chen, F., Wang, Y., Wang, H., Dong, Z., Wang, Y., Zhang, M., Li, J., Shao, S., Yu, C., Huan, Z., and Xu, J. (2019) Flaxseed oil ameliorated high-fat-diet-induced bone loss in rats by promoting osteoblastic function in rat primary osteoblasts. Nutr Metab (Lond) 16, 71

56. Moody, L., Shao, J., Chen, H., and Pan, Y. X. (2019) Maternal Low-Fat Diet Programs the Hepatic Epigenome despite Exposure to an Obesogenic Postnatal Diet. Nutrients 11

57. Allard, J. S., Perez, E., Zou, S., and de Cabo, R. (2009) Dietary activators of Sirt1. Mol Cell Endocrinol 299, 58–63

58. Mercken, E. M., Mitchell, S. J., Martin-Montalvo, A., Minor, R. K., Almeida, M., Gomes, A. P., Scheibye-Knudsen, M., Palacios, H. H., Licata, J. J., Zhang, Y., Becker, K. G., Khraiwesh, H., Gonzalez-Reyes, J. A., Villalba, J. M., Baur, J. A., Elliott, P., Westphal, C., Vlasuk, G. P., Ellis, J. L., Sinclair, D. A., Bernier, M., and de Cabo, R. (2014) SRT2104 extends survival of male mice on a standard diet and preserves bone and muscle mass. Aging Cell 13, 787–796

59. Shaw, R. J., Akufo-Tetteh, E. K., Risk, J. M., Field, J. K., and Liloglou, T. (2006) Methylation enrichment pyrosequencing: combining the specificity of MSP with validation by pyrosequencing. Nucleic Acids Res 34, e78

60. Su, X., Liao, L., Shuai, Y., Jing, H., Liu, S., Zhou, H., Liu, Y., and Jin, Y. (2015) MiR-26a functions oppositely in osteogenic differentiation of BMSCs and ADSCs depending on distinct activation and roles of Wnt and BMP signaling pathway. Cell Death Dis 6, e1851

61. Kim, K., Kim, J. H., Kim, I., Lee, J., Seong, S., Park, Y. W., and Kim, N. (2015) MicroRNA-26a regulates RANKL-induced osteoclast formation. Mol Cells 38, 75–80

62. Krzeszinski, J. Y., Wei, W., Huynh, H., Jin, Z., Wang, X., Chang, T. C., Xie, X. J., He, L., Mangala, L. S., Lopez-Berestein, G., Sood, A. K., Mendell, J. T., and Wan, Y. (2014) miR-34a blocks osteoporosis and bone metastasis by inhibiting osteoclastogenesis and Tgif2. Nature 512, 431–435

63. Wang, H., Xie, Z., Hou, T., Li, Z., Huang, K., Gong, J., Zhou, W., Tang, K., Xu, J., and Dong, S. (2017) MiR-125b Regulates the Osteogenic Differentiation of Human Mesenchymal Stem Cells by Targeting BMPR1b. Cell Physiol Biochem 41, 530–542

64. Mizuno, Y., Yagi, K., Tokuzawa, Y., Kanesaki-Yatsuka, Y., Suda, T., Katagiri, T., Fukuda, T., Maruyama, M., Okuda, A., Amemiya, T., Kondoh, Y., Tashiro, H., and Okazaki, Y. (2008) miR-125b inhibits osteoblastic differentiation by down-regulation of cell proliferation. Biochem Biophys Res Commun 368, 267–272

65. Wang, Z., Xie, Q., Yu, Z., Zhou, H., Huang, Y., Bi, X., Wang, Y., Shi, W., Sun, H., Gu, P., and Fan, X. (2015) A regulatory loop containing miR-26a, GSK3beta and C/EBPalpha regulates the osteogenesis of human adipose-derived mesenchymal stem cells. Sci Rep 5, 15280

66. Goettsch, C., Rauner, M., Pacyna, N., Hempel, U., Bornstein, S. R., and Hofbauer, L. C. (2011) miR-125b regulates calcification of vascular smooth muscle cells. Am J Pathol 179, 1594–1600

67. Lu, Y., Zhao, X., Liu, Q., Li, C., Graves-Deal, R., Cao, Z., Singh, B., Franklin, J. L., Wang, J., Hu, H., Wei, T., Yang, M., Yeatman, T. J., Lee, E., Saito-Diaz, K., Hinger, S., Patton, J. G., Chung, C. H., Emmrich, S., Klusmann, J. H., Fan, D., and Coffey, R. J. (2017) lncRNA MIR100HG-derived miR-100 and miR-125b mediate cetuximab resistance via Wnt/beta-catenin signaling. Nat Med 23, 1331–1341

68. Smith, M. S., and Grove, K. L. (2002) Integration of the regulation of reproductive function and energy balance: lactation as a model. Front Neuroendocrinol 23, 225–256

69. Trayhurn, P., Douglas, J. B., and McGuckin, M. M. (1982) Brown adipose tissue thermogenesis is ‘suppressed’ during lactation in mice. Nature 298, 59–60

70. Rolfe, D. F., and Brown, G. C. (1997) Cellular energy utilization and molecular origin of standard metabolic rate in mammals. Physiol Rev 77, 731–758

71. Clowes, E. J., Aherne, F. X., Foxcroft, G. R., and Baracos, V. E. (2003) Selective protein loss in lactating sows is associated with reduced litter growth and ovarian function. J Anim Sci 81, 753–764

72. Kovacs, C. S. (2016) Maternal Mineral and Bone Metabolism During Pregnancy, Lactation, and Post-Weaning Recovery. Physiol Rev 96, 449–547

73. Wendelboe, M. H., Thomsen, J. S., Henriksen, K., Vegger, J. B., and Bruel, A. (2016) Zoledronate prevents lactation induced bone loss and results in additional post-lactation bone mass in mice. Bone 87, 27–36

74. Woodrow, J. P., Sharpe, C. J., Fudge, N. J., Hoff, A. O., Gagel, R. F., and Kovacs, C. S. (2006) Calcitonin plays a critical role in regulating skeletal mineral metabolism during lactation. Endocrinology 147, 4010–4021

75. Conigrave, A. D., Brown, E. M., and Rizzoli, R. (2008) Dietary protein and bone health: roles of amino acid-sensing receptors in the control of calcium metabolism and bone homeostasis. Annu Rev Nutr 28, 131–155

76. Wysolmerski, J. J. (2002) The evolutionary origins of maternal calcium and bone metabolism during lactation. J Mammary Gland Biol Neoplasia 7, 267–276

77. Kovacs, C. S. (2012) The role of vitamin D in pregnancy and lactation: insights from animal models and clinical studies. Annu Rev Nutr 32, 97–123

78. Ardeshirpour, L., Dumitru, C., Dann, P., Sterpka, J., VanHouten, J., Kim, W., Kostenuik, P., and Wysolmerski, J. (2015) OPG Treatment Prevents Bone Loss During Lactation But Does Not Affect Milk Production or Maternal Calcium Metabolism. Endocrinology 156, 2762–2773

79. Elefteriou, F., Benson, M. D., Sowa, H., Starbuck, M., Liu, X., Ron, D., Parada, L. F., and Karsenty, G. (2006) ATF4 mediation of NF1 functions in osteoblast reveals a nutritional basis for congenital skeletal dysplasiae. Cell Metab 4, 441–451

80. Cunha, M. C., Lima Fda, S., Vinolo, M. A., Hastreiter, A., Curi, R., Borelli, P., and Fock, R. A. (2013) Protein malnutrition induces bone marrow mesenchymal stem cells commitment to adipogenic differentiation leading to hematopoietic failure. PLoS One 8, e58872

81. Ammann, P., Zacchetti, G., Gasser, J. A., Lavet, C., and Rizzoli, R. (2015) Protein malnutrition attenuates bone anabolic response to PTH in female rats. Endocrinology 156, 419–428

82. MacDonell, R., Hamrick, M. W., and Isales, C. M. (2016) Protein/amino-acid modulation of bone cell function. Bonekey Rep 5, 827

83. Long, F. (2011) Building strong bones: molecular regulation of the osteoblast lineage. Nat Rev Mol Cell Biol 13, 27–38

84. Wang, H., Hu, Z., Wu, J., Mei, Y., Zhang, Q., Zhang, H., Miao, D., and Sun, W. (2019) Sirt1 Promotes Osteogenic Differentiation and Increases Alveolar Bone Mass via Bmi1 Activation in Mice. J Bone Miner Res 34, 1169–1181

85. Li, Y., Fan, L., Hu, J., Zhang, L., Liao, L., Liu, S., Wu, D., Yang, P., Shen, L., Chen, J., and Jin, Y. (2015) MiR-26a Rescues Bone Regeneration Deficiency of Mesenchymal Stem Cells Derived From Osteoporotic Mice. Mol Ther 23, 1349–1357

86. Krishnan, V., Bryant, H. U., and Macdougald, O. A. (2006) Regulation of bone mass by Wnt signaling. J Clin Invest 116, 1202–1209

87. Sims, N. A. (2016) Cell-specific paracrine actions of IL-6 family cytokines from bone, marrow and muscle that control bone formation and resorption. Int J Biochem Cell Biol 79, 14–23

